# Sex-specific modulation of the medial prefrontal cortex by glutamatergic median raphe neurons

**DOI:** 10.1101/2023.08.30.555555

**Authors:** Stuart A. Collins, Hannah E. Stinson, Amanda Himes, Ipe Ninan

## Abstract

The current understanding of the neuromodulatory role of the median raphe nucleus (MRN) is primarily based on its putative serotonergic output. However, a significant proportion of raphe neurons are glutamatergic. The present study investigated how glutamatergic MRN input modulates the medial prefrontal cortex (mPFC), a critical component of the fear circuitry. Our studies show that VGLUT3-expressing MRN neurons modulate VGLUT3- and somatostatin-expressing neurons in the mPFC. Consistent with this modulation of mPFC GABAergic neurons, activation of MRN (VGLUT3) neurons suppresses mPFC pyramidal neuron activity and attenuates fear memory in female but not male mice. In agreement with these female-specific effects, we observed sex differences in glutamatergic transmission onto MRN (VGLUT3) neurons and mPFC (VGLUT3) neuron-mediated dual release of glutamate and GABA. Thus, our results demonstrate a cell type-specific modulation of the mPFC by MRN (VGLUT3) neurons and reveal a sex-specific role of this neuromodulation in mPFC synaptic plasticity and fear memory.

## Introduction

The medial prefrontal cortex (mPFC) plays an essential role in regulating fear memory (*1*). While the dorsal mPFC (prelimbic, PL-mPFC) is involved in the expression of fear memory, the ventral mPFC (infralimbic, IL-mPFC) plays a crucial role in fear extinction, an inhibitory mechanism that suppresses fear memory (*1–2*). Although fear memory associated with a potential threat is necessary for survival, a diminished ability to regulate fear memory in a safe environment is implicated in anxiety and psychological trauma-related disorders (*3–5*). Therefore, therapeutic interventions to attenuate fear memory might benefit the management of these disorders. An evidence-based approach to suppressing fear memory requires a better understanding of the neuromodulatory mechanisms in the fear circuitry. Activity in the median raphe nucleus (MRN), which projects to the mPFC, has been implicated in fear memory and anxiety-related behaviors (*6–13*). However, the cellular specificity and synaptic mechanisms underlying the MRN modulation of the mPFC are unclear.

In addition to GABAergic and 5-HTergic neurons, the MRN contains neurons expressing vesicular glutamate transporters VGLUT2 and VGLUT3 (*14–17*). VGLUT2 neurons do not express VGLUT3, GABAergic or 5-HTergic markers (*15*). Approximately 70% of MRN projections to the mPFC are VGLUT3-positive (*10*). Despite the robust glutamatergic nature of the MRN input to the mPFC, activation of the MRN enhances inhibitory activity in the mPFC (*18*). Therefore, it is possible that MRN (VGLUT3) neurons inhibit mPFC pyramidal neurons by activating local GABAergic neurons. A recent study showed that raphe neurons innervate GABAergic neurons in the mPFC (*19*). This purported suppression of pyramidal neuron activity might affect fear memory. Using a cell-type specific expression of channelrhodopsin (ChR2), we show that MRN (VGLUT3) neurons modulate VGLUT3- and somatostatin (SST)-expressing neurons in the mPFC via glutamatergic transmission. Furthermore, activation of MRN (VGLUT3) axonal terminals induces a long-lasting enhancement of GABAergic transmission in the female mPFC. This activation of MRN (VGLUT3) axonal terminals also suppresses excitatory synaptic transmission in the female PL-mPFC. Consistent with the sex difference in MRN (VGLUT3) neuron-mediated plasticity in the mPFC, chemogenetic activation of MRN (VGLUT3) neurons suppresses fear memory in female mice but not male mice. In agreement with these sex-specific effects in mPFC plasticity and fear memory suppression, we observed sex difference in glutamatergic input to MRN (VGLUT3) neurons and the synaptic output of mPFC (VGLUT3) neurons, a primary synaptic target of MRN (VGLUT3) neurons.

## Results

### MRN (VGLUT3) neurons activate VGLUT3 and SST neurons in the mPFC

First, we determined whether MRN (VGLUT3) neurons innervate the mPFC in a cell type-specific manner. Recent studies show colocalization of VGLUT3-expressing raphe axonal boutons with cortical VGLUT3 neurons, most of which are cholecystokinin (CCK)-expressing GABAergic neurons that release both glutamate and GABA (*20–23*). We examined MRN (VGLUT3) neuron-mediated transmission in mPFC (VGLUT3) neurons. Light-evoked currents were recorded at −60mV in tdTomato-expressing mPFC (VGLUT3) neurons from VGLUT3-tdTomato mice (generated by crossing B6;129S-*Slc17a8^tm1.1(cre)Hze^*/J mice with B6.Cg-Gt(ROSA)26Sortm14(CAG-tdTomato)Hze/J mice) that received pAAV-EF1a-double floxed-hChR2(H134R)-mCherry/EYFP-WPRE-HGHpA into the MRN. Of the 201 mPFC (VGLUT3) neurons (116 neurons from 11 female+85 neurons from 7 male mice) recorded, 21 (12 from female mice+9 from male mice) neurons showed synaptic currents in response to light activation of MRN (VGLUT3) terminals (mean amplitude: 28.16±10.09pA) (Figure 1B). We examined whether MRN (VGLUT3) neuron-mediated transmission in mPFC (VGLUT3) neurons is monosynaptic by studying the effect of 4-aminopyridine (4-AP, 100μM), a blocker of voltage-gated potassium channels, on tetrodotoxin (TTX, 1μM)-induced suppression of light-evoked currents in mPFC (VGLUT3) neurons (Figure 1B, C) (*24*). Only 4 out of 9 neurons showed 4-AP-induced rescue of EPSCs, suggesting that MRN (VGLUT3) neurons activate mPFC (VGLUT3) neurons via both monosynaptic and polysynaptic connections. The average delay between the onset of the light stimulus and synaptic currents before the application of TTX were 1.91±0.13ms and 8.21±0.4ms for monosynaptic (4-AP responsive) and polysynaptic (4-AP unresponsive) currents, respectively. The currents in the presence of both TTX and 4-AP were blocked by DNQX (10 μM), an AMPA receptor blocker, confirming the glutamatergic nature of this synaptic transmission (Figure 1B). Apart from MRN (VGLUT3) neuron-mediated glutamatergic transmission in mPFC (VGLUT3) neurons, we observed that 11 out of 56 presumed SST neurons (13 mice) showed synaptic currents in response to the light activation (mean amplitude: 21.03±4.02pA). These SST neurons were identified based on their active and passive membrane properties (*25*). To confirm MRN (VGLUT3) neuron-mediated transmission in SST neurons in the mPFC, we studied light-evoked currents in EGFP-expressing SST neurons from MRN (VGLUT3)-ChR2-GIN mice. To generate MRN (VGLUT3)-ChR2-GIN mice, we first crossed B6;129S-*Slc17a8^tm1.1(cre)Hze^*/J mice with FVB-Tg(GadGFP)45704Swn/J (GIN) mice (*26–27*). Then, the progeny expressing both Cre and EGFP were injected with pAAV-EF1a-double floxed-hChR2(H134R)-mCherry-WPRE-HGHpA into the MRN. We observed that 7 out of 119 EGFP-expressing SST neurons (12 female+9 male mice) exhibited synaptic currents in response to a light activation (mean amplitude: 18±3.09pA) and that these currents were blocked by DNQX confirming the glutamatergic nature of this synaptic transmission (Figure 1D). The average delay between the onset of light stimulus and synaptic response was 5.3±0.8ms, suggesting that the MRN (VGLUT3) neuron modulation of mPFC (SST) neurons is not monosynaptic. Consistently, bath application of 4-AP failed to rescue TTX-induced blockade of MRN (VGLUT3) neuron-mediated currents in mPFC (SST) neurons (Figure 1D). In contrast to MRN (VGLUT3)-mediated synaptic transmission in SST and VGLUT3 neurons in the mPFC, we did not observe light-evoked synaptic responses in EGFP-expressing parvalbumin (PV) neurons (n=110 neurons/4 female+3male mice) of MRN (VGLUT3)-ChR2-G42 mice [mice expressing ChR2 in MRN (VGLUT3) neurons and EGFP in PV neurons] (*28*). MRN (VGLUT3)-ChR2-G42 mice were generated using the same breeding approach used for MRN (VGLUT3)-ChR2-GIN mice. Similar to the results in PV neurons, none of the recorded pyramidal neurons (134 neurons/22 female+22 male mice) showed a synaptic response to the 5ms light pulse activation. However, a 10Hz light stimulation for 60s (each pulse of 2ms duration) produced a hyperpolarization response in 15 (5 from female mice and 10 from male mice) out of 82 layer 5 pyramidal cells in MRN (VGLUT3)-ChR2 mice (20 female +20 male mice) (Figure 1F). These results suggest that MRN (VGLUT3) neurons exert a GABAergic inhibition of pyramidal neurons in the mPFC by activating local VGLUT3 and SST neurons. The MRN (VGLUT3) neuron-mediated monosynaptic activation of mPFC (VGLUT3) neurons might be responsible for polysynaptic glutamatergic responses in SST and VGLUT3 neurons in the mPFC.

**Figure 1.**
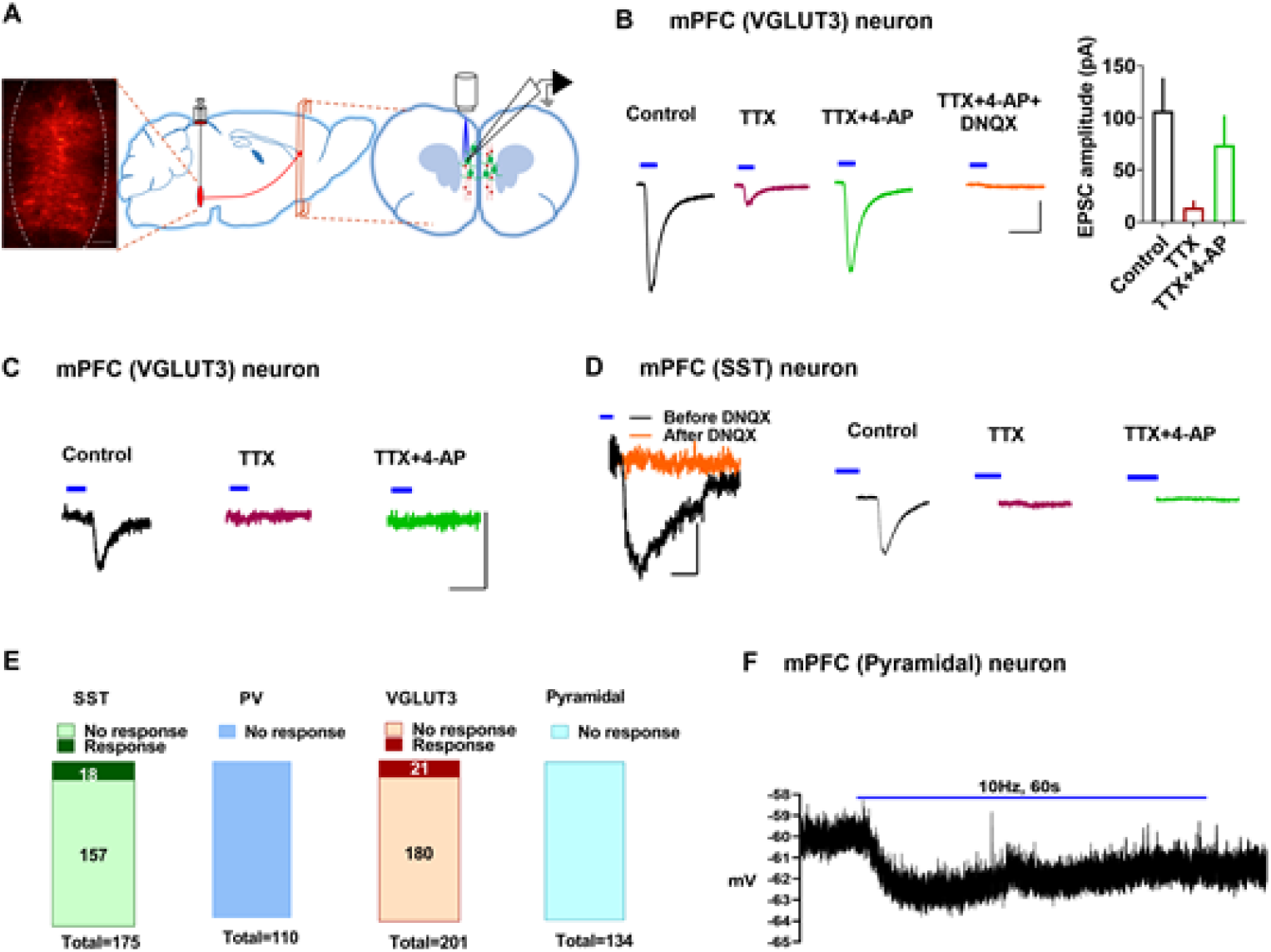
MRN (VGLUT3) neuron modulation of the mPFC. A) Stereotactic injection of pAAV-EF1a-double floxed-hChR2(H134R)-mCherry-WPRE-HGHpA nto the MRN of VGLUT3 Cre mice. The left panel shows an example of mC he rry-expres sing MRN (VGLUT3) neurons 21 days after the injection. Scale 100 pM The right panel shows a schematic of m Che rry-expressing MRN (VGLUT3) axonal fibers (shown in red) in the mPFC and whole-cell recording of mPFC neurons (shown in green) in the brain slice preparation B) Monosynaptic MRN (VGLUT3-mPFC (VGLUT3) glutamatergic transmission. Effect of 4-AP (100 pM) on TTX (1 pM)-induced suppression of light-evoked currents. DNQX (10 pM), an AMPA receptor antagonist blocked MRN (VGLUT3) neuron-mediated currents in mPFC (VGLUT3) neurons. Scale: 10 ms/50 pA. The right panel shows the mean EPSC amplitude in 4 mPFC (VGLUT3) neurons before TTX (control) in the presence of TTX and TTX+4-AP. C) Polysynaptic MRN (VGLUT3) neuron-mediated transmission in mPFC (VGLUT3) neurons. In some mPFC (VGLUT3) neurons, 4-AP failed to rescue the TTX-induced block of EPSCs. Scale: 10 ms/25 pA. D) Left panel shows an example of MRN (VGLUT3) neuron-mediated glutamatergic current in an mPFC (SST) neuron from MRN (VGLUT3)-ChR2-GIN mice. DNQX blocked VGLUT3 neuron-mediated currents in SST neurons. Scale 10 ms/10 pA. 4-AP failed to rescue the TTX block of MRN (VGLUT3) neuron-mediated currents in SST neurons. Scale: 10 ms/50 pA E) Summary of results from experiments to examine MRN (VGLUT3) neuron-mediated transmission in SST PV. VGLUT3, and pyramidal neurons in the mPFC. SST and VGLUT3 neurons but not PV or pyramidal neurons, showed synaptic currents in response to light activation of MRN (VGLUT3) terminals in the mPFC. F) An example trace showing hyperpolarization in pyramidal neurons in response to a 10 Hz light stimulation (2 ms pulse) for GO s. Blue lines indicate the application of light (470 nm).

### MRN (VGLUT3) neuron-mediated long-lasting enhancement of GABAergic transmission in the PL-mPFC pyramidal neurons of female but not male mice

Based on our data showing MRN (VGLUT3) neuron-mediated glutamatergic currents in VGLUT3 and SST neurons in the mPFC, we asked whether activity in MRN (VGLUT3) axonal terminals affects GABAergic transmission in layer 5 pyramidal neurons. To study the effect of light-induced activation of MRN (VGLUT3) terminals in the mPFC, we used MRN (VGLUT3)-ChR2 mice, as described above. As controls, we used MRN-EGFP mice which received pAAV-hSyn-EGFP injection into the MRN. MRN (VGLUT3) terminals were activated by a 10 Hz light stimulation for 60s (each pulse of 2ms duration). Spontaneous inhibitory post-synaptic currents (sIPSCs) were recorded at 0 mV before and after the 10 Hz stimulation. This 10 Hz stimulation produced a long-lasting enhancement of both the frequency and amplitude of sIPSCs in the PL-mPFC pyramidal neurons of female MRN (VGLUT3)-ChR2 mice but not MRN-EGFP mice (Figure 2A). However, this GABAergic plasticity in PL-mPFC pyramidal neurons was absent in male MRN (VGLUT3)-ChR2 mice (Figure 2B). These results demonstrate a sex difference in MRN (VGLUT3) neuron-mediated GABAergic plasticity in PL-mPFC pyramidal neurons.

**Figure 2.**
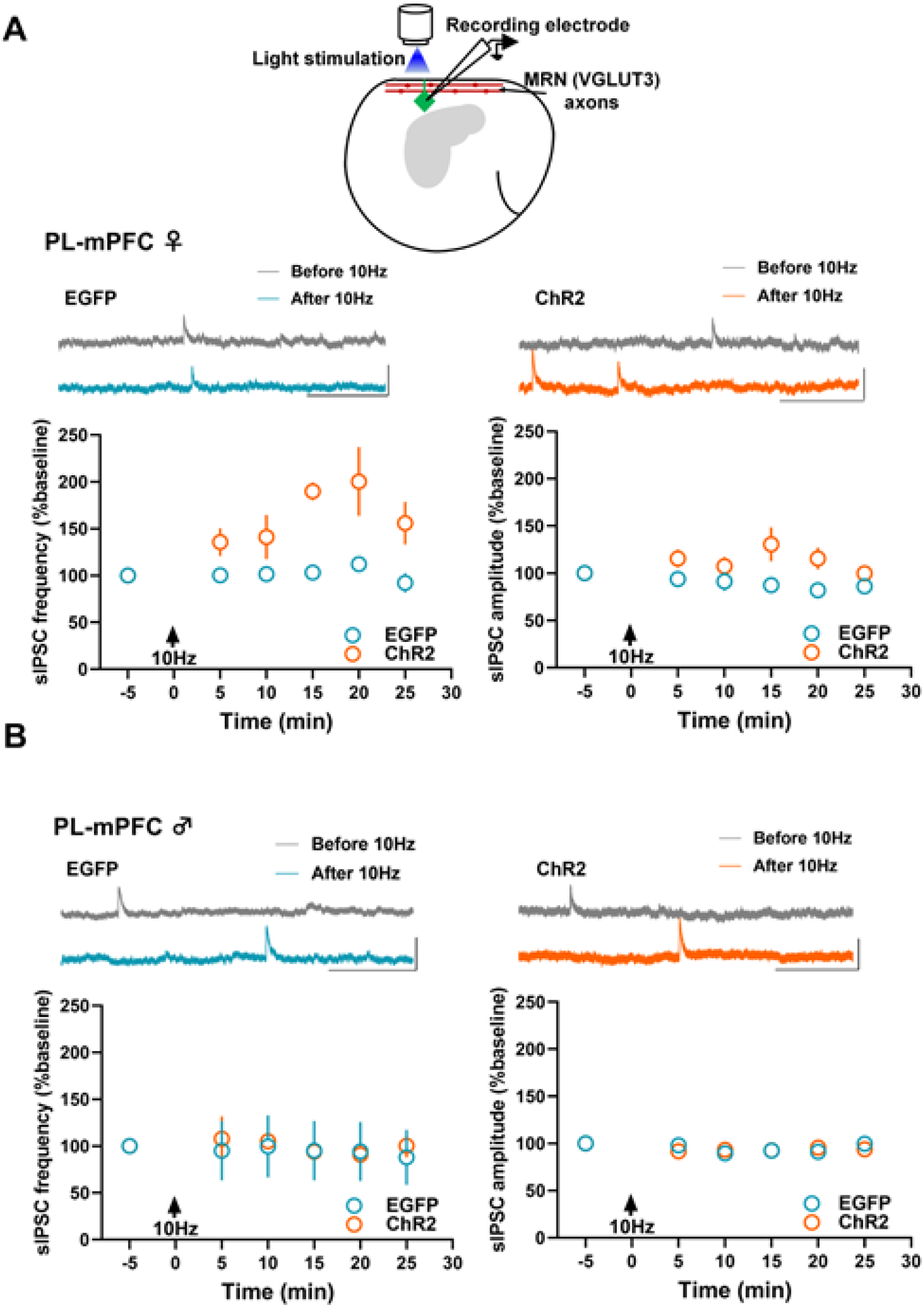
Activation of MRN (VGLUT3) terminals enhances GABAergic transmission in the PL-mPFC pyramidal neurons of female but not male mice. A) Upper panel shows a schematic of whole-cell recording of pyramidal neurons and the application of 10Hz light stimulation to activate MRN (VGLUT3) terminals A 10 Hz light stimulation (pulse duration: 2 ms) for 60 s enhanced sIPSC frequency (F_1.17_=16.9, P<0.001) and amplitude (F_1.17_=4.7, P=0.046) in PL-mPFC pyramidal neurons of female MRN (VGLUT3)-ChR2 group (10 neurons/5 mice) compared to MRN-EGFP group (9 neurons/5 mice). B) PL-mPFC pyramidal neurons in male MRN (VGLUT3)-ChR2 group (9 neurons/5 mice) did not show a significant change in either frequency (F_1.16_=0.05, P=0.83) or amplitude (F_1,16_=0.08, P=0.77) of sIPSCs as compared to MRN-EGFP group (9 neurons/5 mice). The upper panel shows examples traces. Scale: 100 pA/250 ms.

Next, we examined whether a similar MRN (VGLUT3) neuron-mediated GABAergic plasticity occurs in the IL-mPFC. Although the 10 Hz light stimulation showed a late increase in sIPSC frequency in the female MRN (VGLUT3)-ChR2 group compared to the MRN-EGFP group, we did not observe any change in sIPSC amplitude in female mice after the 10Hz light stimulation (Figure 3A). We did not observe any change in sIPSC amplitude or frequency in the male MRN (VGLUT3)-ChR2 group compared to the MRN-EGFP group (Figure 3B). These results show that activation of MRN (VGLUT3) terminals produces a sex- and region-specific GABAergic plasticity in mPFC pyramidal neurons.

**Figure 3.**
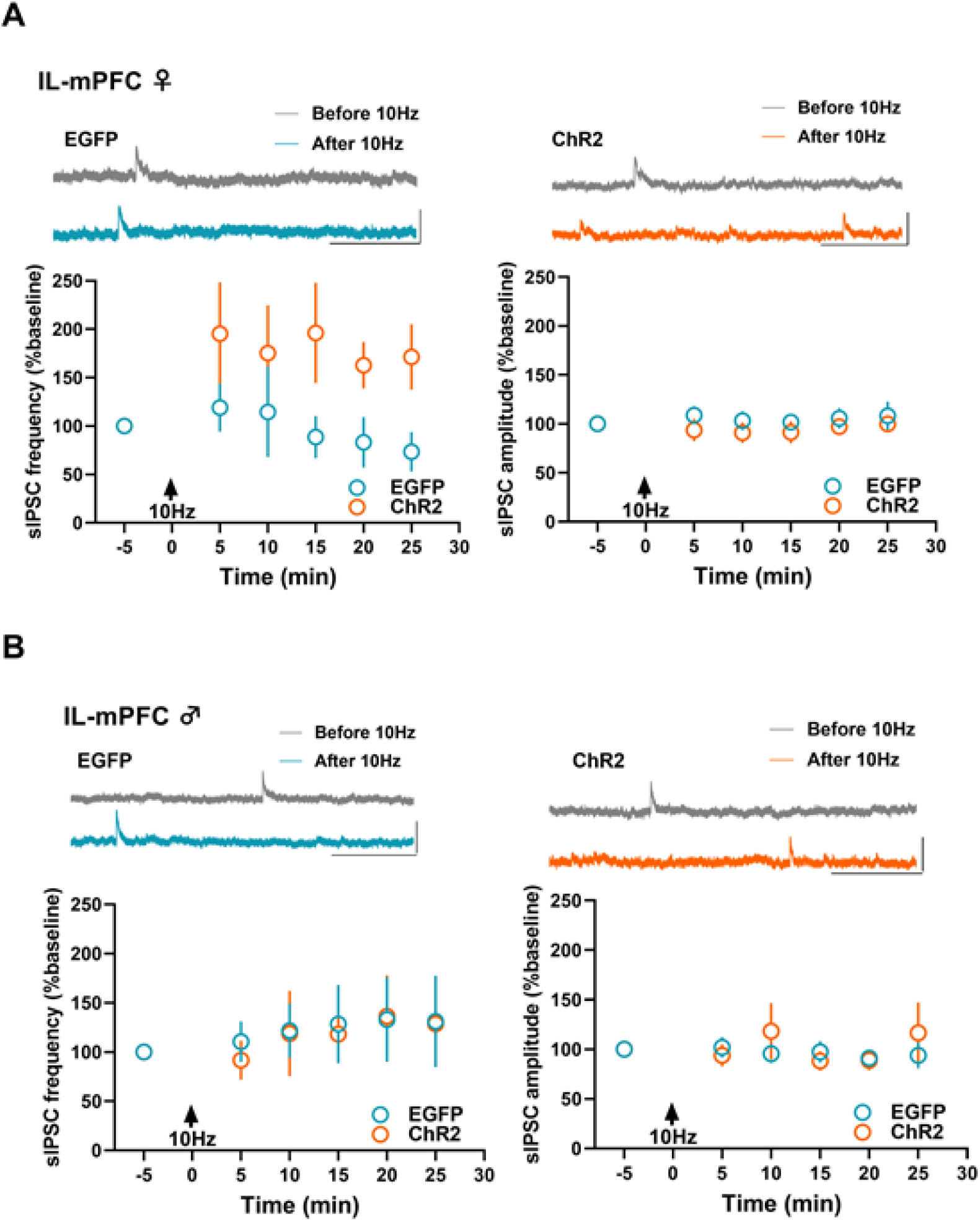
Effect of MRN (VGLUT3) terminal activity on GABAergic transmission in IL-mPFC pyramidal neurons. A) Although 10 Hz light stimulation showed a delayed effect on sIPSC frequency (F_1,19_=4.45, P=0.048, last 15 min), overall frequency (F_1,19_=3.03, P=0.09) and amplitude (F_1,19_=0.92, P=0.34) following the light activation did not reveal a statistically significant effect between female MRN (VGLUT3)-ChR2 group (12 neurons/5 mice) and MRN-EGFP group (9 neurons/5 mice). B) The 10 Hz light activation did not affect either frequency (F_1,16_=0.02, P=0.88) or amplitude (F_1,16_=0.07, P=0.78) of sIPSCs in the IL-mPFC pyramidal neurons of male MRN (VGLUT3)-ChR2 group (9 neurons/5 mice) compared to MRN-EGFP group (9 neurons/5 mice). The upper panel shows examples traces. Scale: 100 pA/250 ms.

### MRN (VGLUT3) neuron-mediated suppression of excitatory synaptic transmission in the PL-mPFC of female but not male mice

Given the long-lasting enhancement of GABAergic transmission in the PL-mPFC pyramidal neurons of female mice following the activation of MRN (VGLUT3) terminals, we asked whether this activity affects excitatory synaptic transmission in mPFC pyramidal neurons. We measured excitatory post-synaptic potential (EPSP) slope in PL-mPFC and IL-mPFC layer 5 pyramidal neurons before and after a 10Hz light stimulation. We observed a long-lasting depression of EPSP slope in the PL-mPFC pyramidal neurons of female MRN (VGLUT3)-ChR2 mice (Figure 4A). However, this 10Hz stimulation did not produce a statistically significant effect in male PL-mPFC pyramidal neurons or female IL-mPFC pyramidal neurons (Figures 4B-C). Interestingly, we observed a delayed suppression of EPSP slope in male IL-mPFC pyramidal neurons (Figure 4D). These results suggest that MRN (VGLUT3) neuron activity selectively suppresses excitatory synaptic activity in a region- and sex-specific manner.

**Figure 4.**
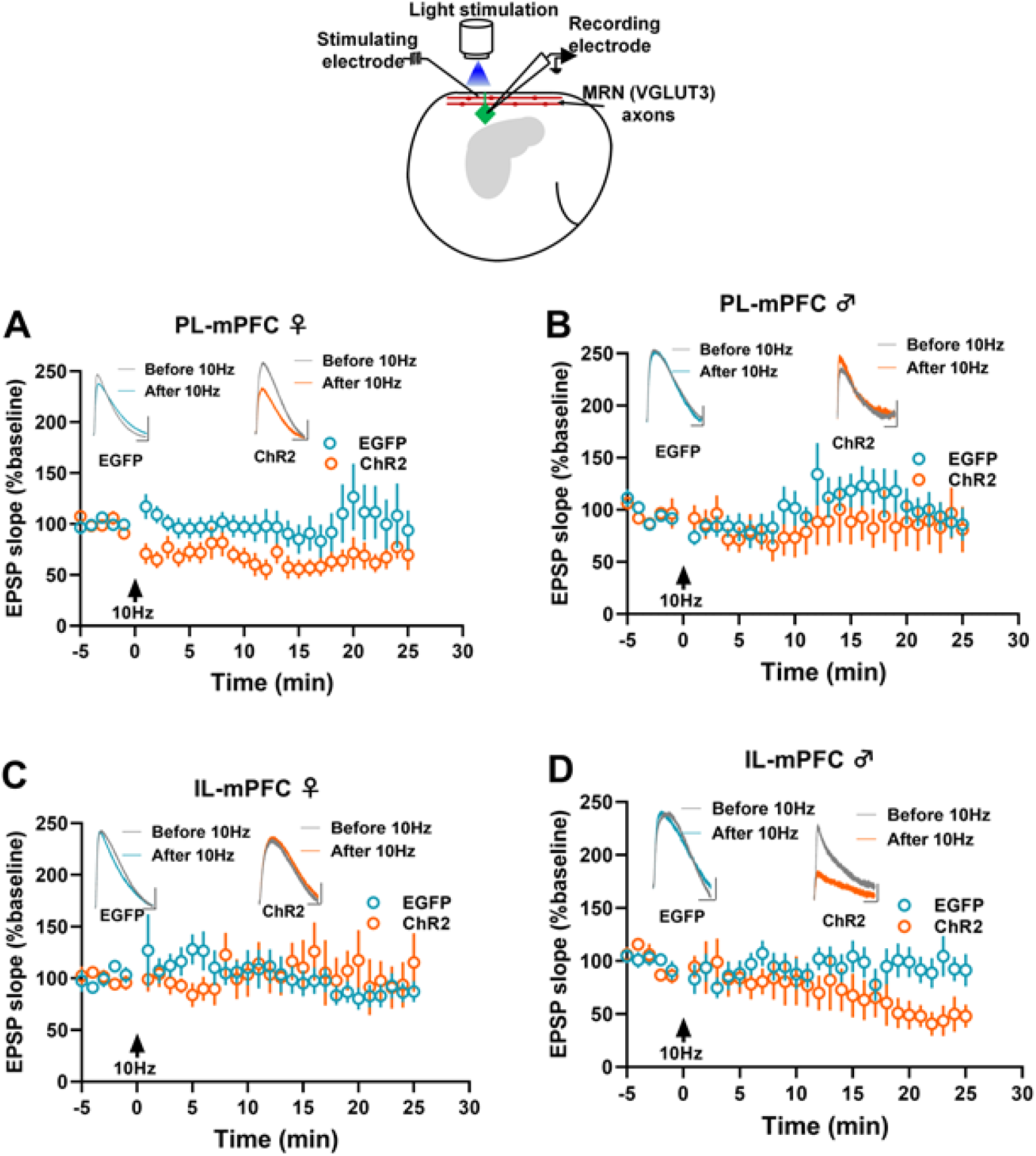
Activation of MRN (VGLUT3) terminals suppresses excitatory synaptic transmission in the PL-mPFC pyramidal neurons of female mice. The upper panel shows a schematic of whole-cell recording of EPSPs in layer 5 pyramidal neurons by field stimulation of layer 2/3 and applying 10Hz light stimulation to activate MRN (VGLUT3) terminals. A) A 10 Hz light stimulation for 60 s showed suppression of EPSP slope (F_1,19_=5.6, P=0.02) in the PL-mPFC pyramidal neurons of female MRN (VGLUT3)-ChR2 group (12 neurons/6mice) compared to MRN-EGFP group (9 neurons/5 mice). B) Light stimulation at 10 Hz did not affect EPSP slope (F_1,17_=0.67, P=0.4) in the PL-mPFC pyramidal neurons of male MRN (VGLUT3)-ChR2 group (10 neurons/5mice) compared to MRN-EGFP group (9 neurons/5 mice). C) The IL-mPFC pyramidal neurons of the female MRN (VGLUT3)-ChR2 group (9 neurons/6mice)did not show a change in EPSP slope (F_1,16_=0.4, P=0.85) compared to MRN-EGFP group (9 neurons/5 mice) after the 10 Hz light stimulation D) Although the IL-mPFC pyramidal neurons of male MRN (VGLUT3)-ChR2 group (10 neurons/5mice) showed a delayed suppression of EPSP slope after the light stimulation (last 5 min, F_1,18_=6.36, P=0.02), a comparison of overall data in MRN (VGLUT3)-ChR2 and MRN-EGFP (10 neurons/5 mice) groups after the light activation did not show a statistically significant effect (F_1,18_=1.5, P=0 23) The inset shows example traces. Scale: 1 mV/25 ms

### Activation of MRN (VGLUT3) neurons does not affect membrane properties of mPFC pyramidal neurons

Although the light activation of MRN (VGLUT3) terminals produced sex- and region-specific changes in GABAergic and glutamatergic transmission in the mPFC, our analysis of passive and active membrane properties in PL-mPFC and IL-mPFC layer 5 pyramidal neurons did not show an effect of MRN (VGLUT3) neuronal activity on input resistance, time constant, capacitance, resting membrane potential or frequency of action potentials evoked by current injection (Figures S1-S4). These results suggest that the activation of MRN (VGLUT3) neurons selectively affects the synaptic properties but not the membrane properties of mPFC pyramidal neurons.

### Activation of MRN (VGLUT3) neurons suppresses fear memory in female but not male mice

Our results show that activity in MRN (VGLUT3) neurons produces a robust and long-lasting increase in GABAergic transmission and suppression of excitatory synaptic transmission in the PL-mPFC of female mice. Given the involvement of the mPFC in regulating fear memory and fear extinction, we examined whether activation of MRN (VGLUT3) neurons affects fear memory or extinction. We studied fear memory and extinction in mice expressing hM3D(Gq) in MRN (VGLUT3) neurons [MRN (VGLUT3)-hM3D(Gq) mice, B6;129S-*Slc17a8^tm1.1(cre)Hze^*/J mice that received pAAV-hSyn-DIO-hM3D(Gq)-mCherry injection into the MRN]. Mice were used 3 weeks after the surgery. On day 1 of the behavioral experiments, mice underwent fear conditioning. On day 2, the fear-conditioned mice received either saline or clozapine N-oxide (CNO, 1 mg/kg), followed by fear extinction training 1 hr later. On day 3, all mice were tested for extinction memory. We did not observe a difference in freezing between the CNO and saline groups on day 3 in both males and females (Figures 5C, D), suggesting a lack of effect of MRN (VGLUT3) neuron activation on fear extinction. However, a comparison of the first three trials during the extinction training on day 2 revealed that freezing is significantly lower in the female CNO group compared to the female saline group (Figure 5C). This effect of MRN (VGLUT3) neuron activation on fear memory was lacking in male mice (Figure 5D), suggesting that activation of MRN (VGLUT3) neurons suppresses fear memory only in females. In control experiments, CNO (1 mg/kg) did not affect fear memory or extinction in mice that received a pAAV-hSyn-EGFP injection into the MRN, suggesting the absence of non-specific effects of CNO on fear memory or extinction (Figure S5).

**Figure 5.**
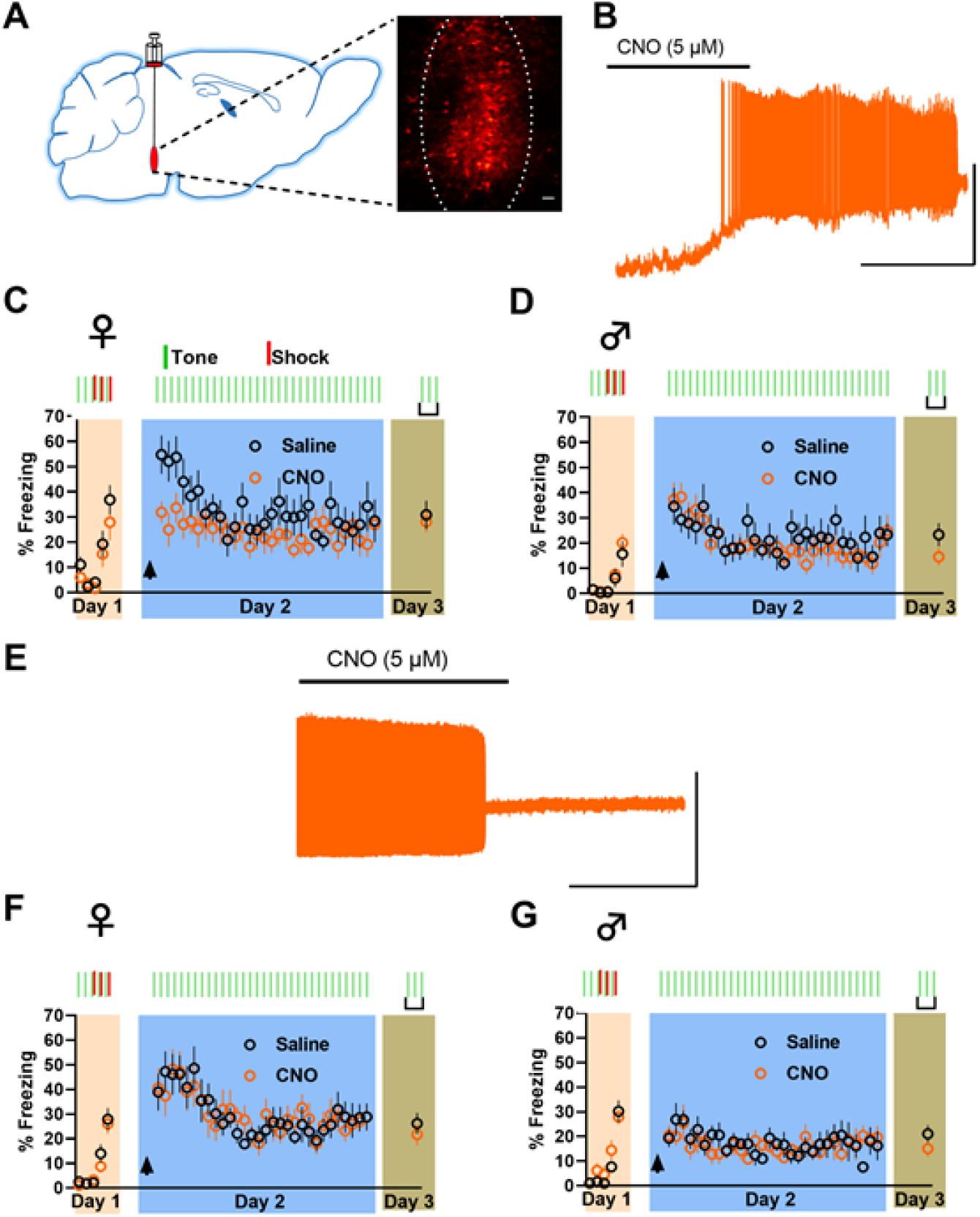
Activation of MRN (VGLUT3) neurons suppresses fear memory in female mice. A) A schematic of viral injection into the MRN and an example of brain slice from VGLUT3 Cre mice showing mCherry-expressing MRN (VGLUT3) neurons 3 weeks after the injection of pAAV-hSyn-DIO-hM3D(Gq)-mCherry. B) An example trace showing the induction of action potentials in a hM3D(Gq)- expressing MRN (VGLUT3) neuron after the perfusion of CNO (5 pM). Scale: 50 mV/100 s. C) Intraperitoneal injection of CNO (1 mg/kg, n=10) but not saline (n=9) 1 hr before extinction training caused a decrease in freezing during the early trials of extinction training (initial 3 trials) in female mice, suggesting suppression of fear memory (F_1,17_=7.8, P=0.01). D) Freezing in CNO (n=10)- and saline (n= 10 (-treated male groups did not show a statistically significant effect (F_1,18_=0.25, P=0.62). Arrow indicates the injection of saline/CNO. E) An example trace showing the suppression of action potentials in a spontaneously firing hM4D(Gi)-expressing MRN (VGLUT3) neuron after the perfusion of CNO (5 pM). Scale: 50 mV/100 s. F) CNO (n=14) and saline (n=13) treatment in hM4D(Gi)- expressing female mice did not show a significant difference in freezing (F_1,25_=0.001, P=0.99) G) CNO (n=9) and saline (n=9) treatment in hM4D(Gi)- expressing male mice did not show a significant difference in freezing (F_1,16_=0.12, P=0.73) Arrow indicates the injection of saline/CNO.

Next, we tested whether suppression of activity in MRN (VGLUT3) neurons affects fear memory or extinction. We studied fear memory and extinction in mice expressing hM4D(Gi) in MRN (VGLUT3) neurons [MRN (VGLUT3)-hM4D(Gi) mice, B6;129S-*Slc17a8^tm1.1(cre)Hze^*/J mice that received pAAV-hSyn-DIO-hM4D(Gi)-mCherry injection into the MRN]. Neither female nor male mice showed an effect on fear memory or extinction after the chemogenetic suppression of MRN (VGLUT3) neurons (Figure 5F, G). These results suggest that only an increase but not a decrease in the activity of MRN (VGLUT3) neurons affects fear memory in female mice.

### Fear conditioning enhances glutamatergic transmission onto MRN (VGLUT3) neurons in female but not male mice

Since MRN (VGLUT3) neuronal activity affects PL-mPFC synaptic transmission and fear memory in female mice, we examined whether fear conditioning or extinction involves changes in synaptic transmission onto MRN (VGLUT3) neurons. We recorded spontaneous glutamatergic and GABAergic transmission in tdTomato-expressing MRN neurons from control tone-alone (TA), fear-conditioned (FC), and fear-extinguished (FE) VGLUT3-tdTomato mice. Miniature excitatory post-synaptic currents (mEPSCs) and miniature inhibitory post-synaptic currents (mIPSCs) were recorded in VGLUT3 neurons at −50 mV and 0 mV, respectively. Interestingly, we observed an enhanced mEPSC amplitude in the FC female group compared to the TA female group (Figure 6B). However, this increase in mEPSC amplitude was not associated with any significant change in mEPSC frequency. Furthermore, the mEPSC amplitude of the FC group was not significantly different from that of the FE group, suggesting that fear extinction does not reverse fear conditioning-induced increase in AMPA receptor-mediated synaptic transmission in MRN (VGLUT3) neurons. Neither fear conditioning nor extinction affected mEPSCs in male mice (Figure 6C). A comparison of mIPSC frequency and amplitude among the TA, FC, and FE groups did not show a statistically significant effect in female or male mice (Figures 6B, C). These results indicate that fear conditioning involves an enhancement of glutamatergic but not GABAergic transmission in MRN (VGLUT3) neurons and that this experience-dependent plasticity occurs only in female mice.

**Figure 6.**
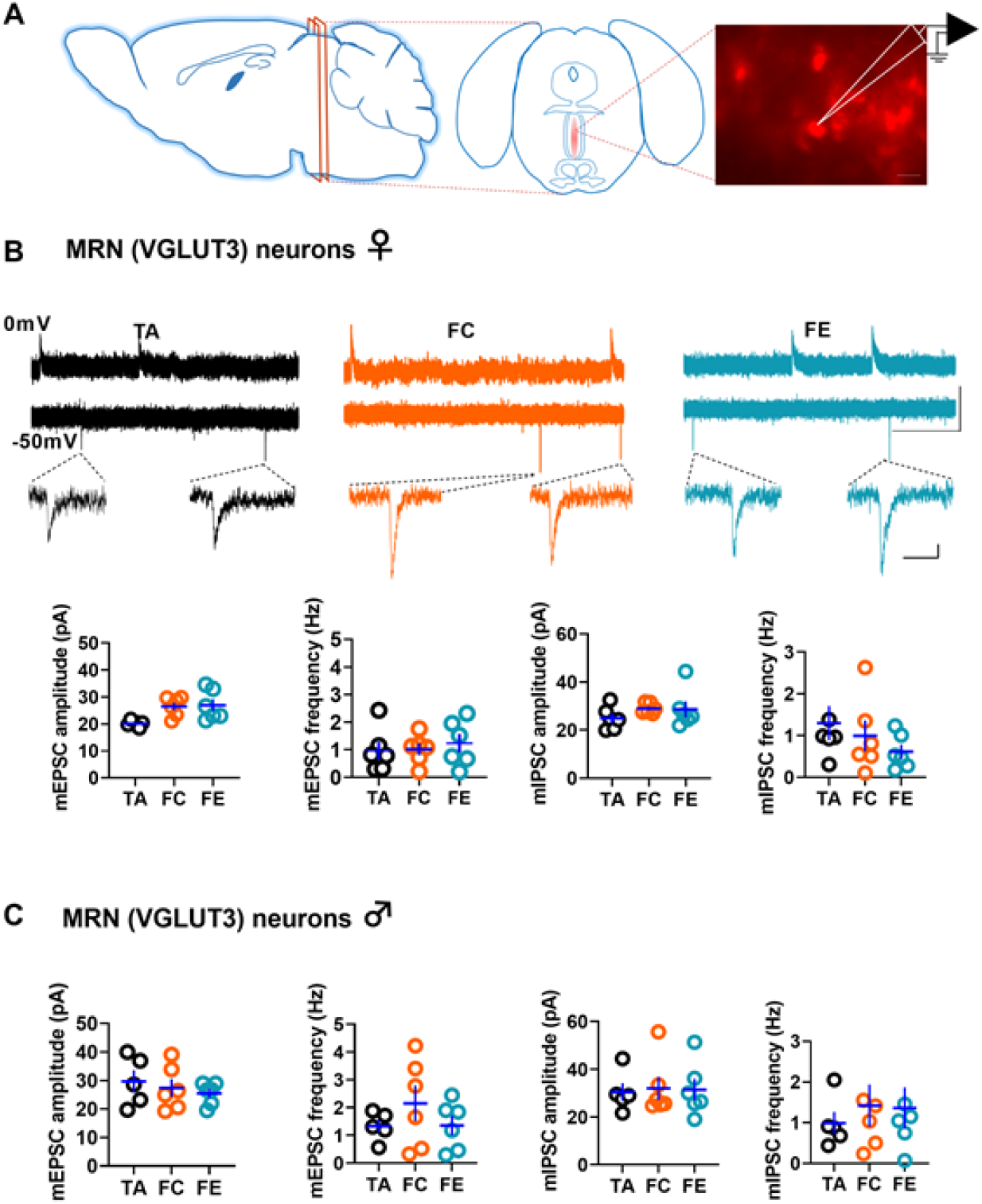
Fear conditioning enhances excitatory synaptic transmission in the MRN (VGLUT3) neurons of female but not male mice. A) A schematic of MRN slice preparation and the recording of tdTomato-expressing MRN (VGLUT3) neurons. B) Comparison of spontaneous excitatory and inhibitory synaptic transmission in MRN (VGLUT3) neurons from female TA (13 neurons/6 mice), FC (13 neurons/6 mice), and FE (16 neurons/6 mice) groups. MRN (VGLUT3) neurons from the female FC group showed a significantly higher mEPSC amplitude than the TA group. Fear extinction did not affect the increased mEPSC amplitude in the FC group (F_2,15_=8.3, P=0.004, TA vs. FC: P=0.006, TA vs. FE: 0.016, FC vs. FE: P=1). None of the other synaptic measurements, i.e., mEPSC frequency (F_2,15_=0.24, P=0.78), mIPSC amplitude (F_2,15_=0.98, P=0.39), and mIPSC frequency (F_2,15_=1.O5, P=0.37) showed statistical significance. The upper panel shows examples of mIPSCs (upper continuous traces) and mEPSCs (lower continuous traces and single events), Scale: 50 pA/500 ms (continuous traces) and 10 pA/10 ms (single events). C) Comparison of spontaneous excitatory and inhibitory synaptic transmission in MRN (VGLUT3) neurons from male TA(13 neurons/5 mice), FC (15 neurons/6 mice), and FE (16 neurons/6 mice) groups. The statistical comparison did not reveal significance in mEPSC amplitude (F_2,14_=0.5, P=0.6), mEPSC frequency (F_2,14_=1.02, P=0.38), mIPSC amplitude (F_2,14_=0.03, P=0.97), and mIPSC frequency (F_2,14_=0.24, P=0.79) among male TA, FC, and FE groups. The statistical comparison showed a significant difference in mEPSC amplitude between female and male TA groups (P=0.02). However, the comparison of mEPSC frequency (P=0.4), mIPSC amplitude (P=0.2), or mIPSC frequency (P=0.6) did not show statistical significance.

Using the control TA groups, we also examined whether the basal glutamatergic or GABAergic transmission in MRN (VGLUT3) neurons exhibits a sex difference. Our analysis showed that the amplitude of mEPSCs is significantly higher in the male TA group compared to the female TA group. However, we did not observe a sex difference in mEPSC frequency, mIPSC amplitude, or mIPSC frequency in TA groups. These results suggest a diminished basal AMPA receptor transmission in the MRN (VGLUT3) neurons of female mice.

We also compared the passive and active membrane properties of VGLUT3 neurons from the TA, FC, and FE groups. Although the female FC group showed an increase in the number of action potentials, this effect did not reach statistical significance (Figure S6A). We did not observe a significant effect of fear conditioning or extinction on membrane properties in male mice (Figure S6B). Also, we did not observe any sex differences in basal passive and active membrane properties except that the female TA group showed a significant depolarization compared to the male TA group (Figures S6A, S6B).

These results suggest that MRN (VGLUT3) neurons in female mice exhibit a diminished glutamatergic input compared to those in male mice, while fear conditioning enhances glutamatergic transmission in the MRN (VGLUT3) neurons of female but not male mice. This sex difference in glutamatergic transmission and experience-dependent glutamatergic plasticity in MRN (VGLUT3) neurons might contribute to female-specific mPFC plasticity and fear memory suppression following the activation of MRN (VGLUT3) neurons.

### Sex difference in PL-mPFC (VGLUT3) neuron-mediated glutamatergic and GABAergic transmission

Since the observed female-specific effect of MRN (VGLUT3) neuronal activation on PL-mPFC synaptic plasticity and fear memory could also involve sex-specific effects within the PL-mPFC, we asked whether PL-mPFC (VGLUT3) neurons modulate other mPFC neurons in a sex-specific manner. We recorded PL-mPFC (VGLUT3) neuron-mediated EPSCs and IPSCs at −20 mV in EGFP-expressing SST neurons and pyramidal neurons from PL-mPFC (VGLUT3)-ChR2-GIN mice, which were generated by injecting pAAV-EF1a-double floxed-hChR2(H134R)-mCherry-WPRE-HGHpA into the PL-mPFC of *Slc17a8^tm1.1(cre)Hze^*/J mice crossed with GIN mice. Although PL-mPFC (VGLUT3) neuron-mediated EPSCs in SST neurons were monosynaptic, we observed both monosynaptic and disynaptic IPSCs, which were blocked by DNQX. The average EPSC and IPSC amplitudes were significantly higher in female mice than in male mice (Figure 7B). We measured the excitatory/inhibitory balance of PL-mPFC (VGLUT3) neuron-mediated transmission in SST neurons by calculating the ratio of EPSC amplitude to IPSC amplitude at a single neuron level (E/I ratio). We did not observe a sex difference in the E/I ratio, suggesting that the overall PL-mPFC (VGLUT3) neuron-mediated transmission in SST neurons is higher in female mice (Figure 7B). Similar to PL-mPFC (VGLUT3) neuron-mediated transmission in SST neurons, PL-mPFC (VGLUT3) neuron-mediated EPSCs in pyramidal neurons were monosynaptic. Also, we observed both monosynaptic and disynaptic IPSCs in pyramidal neurons. Unlike the SST neurons, we did not observe any sex difference in average mPFC (VGLUT3) neuron-mediated EPSC or IPSC amplitude in pyramidal neurons. However, a decreased E/I ratio in female mice compared to male mice suggested that the PL-mPFC (VGLUT3) neuron-mediated inhibition of pyramidal neurons at the single neuron level is higher in female mice (Figure 7C). We also examined PL-mPFC (VGLUT3) neuron-mediated glutamatergic and GABAergic transmission in local PV neurons. Measurement of EPSCs (18.9±8.4pA) and IPSCs (29.2±20.5pA) in EGFP-expressing PV neurons from PL-mPFC (VGLUT3)-ChR2-G42 mice (Figure S7) showed that PL-mPFC (VGLUT3) neuron-mediated transmission in local PV neurons is comparable to that in SST neurons (EPSC=30.4±7.5pA, IPSC=29.6±11.04pA) (Figure 7B) but weaker than that in pyramidal neurons (EPSC=60.9±14.9pA, IPSC=101.3±26.7pA) (Figure 7C). We did not observe sex difference in EPSC amplitude, IPSC amplitude, or E/I ratio in PV neurons. These results suggest that the sex-specific effect of MRN (VGLUT3) neuronal activation on PL-mPFC synaptic plasticity and fear memory suppression might also be due to sex difference in PL-mPFC (VGLUT3) neuron-mediated transmission in local SST and pyramidal neurons.

**Figure 7.**
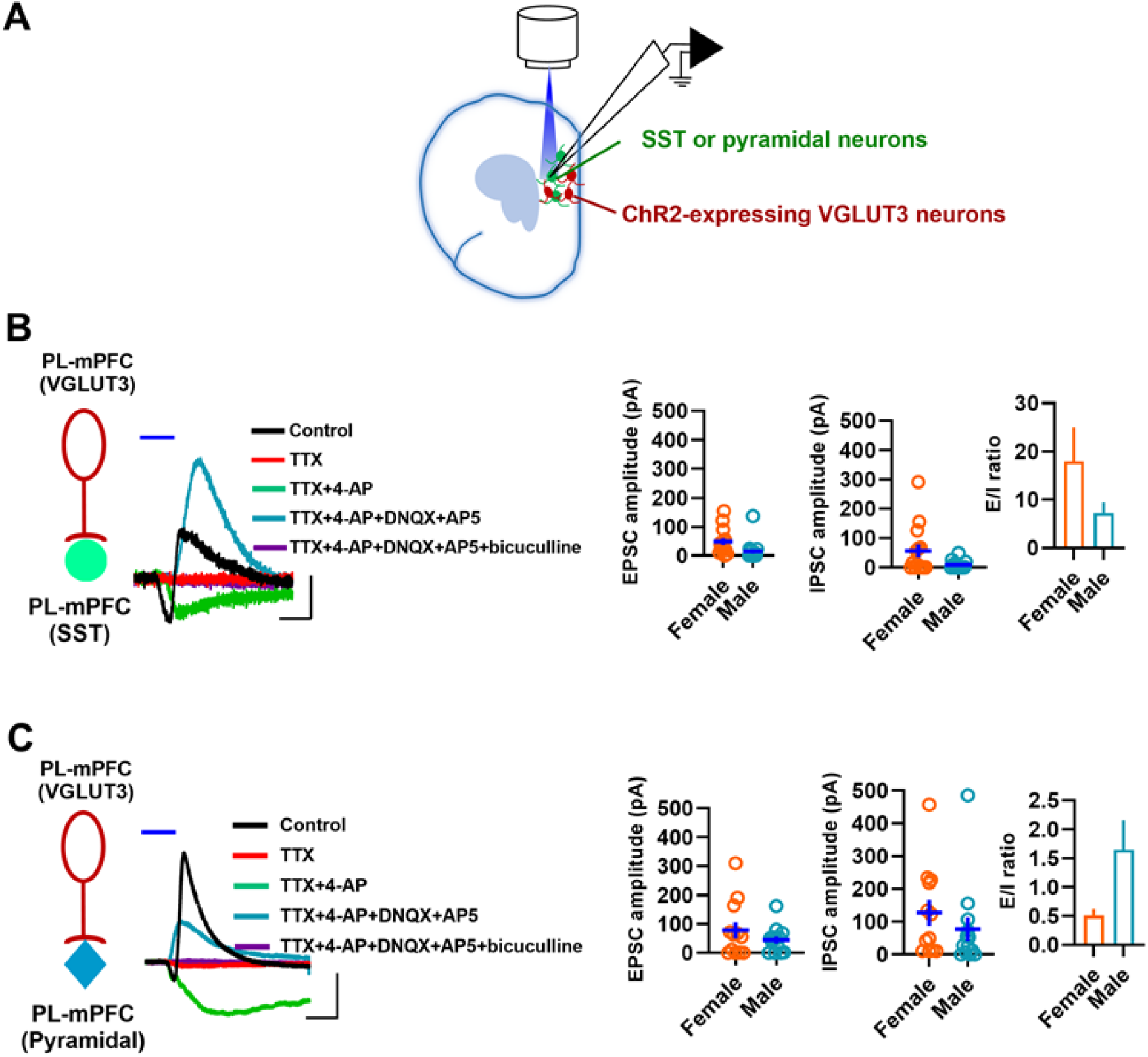
Activation of PL-mPFC (VGLUT3) neurons produces glutamatergic and GABAergic responses in local SST and pyramidal neurons. AJA schematic of light activation of PL-mPFC (VGLUT3) neurons and whole-cell recording in local SST and pyramidal neurons B) EPSC amplitude, IPSC amplitude, and excitatory/inhibitory ratio (Bl ratio) in PL-mPFC (SST) neurons in response to light activation of PL- mPFC (VGLUT3) neurons. We observed monosynaptic EPSCs in SST neurons. While some neurons showed disynaptic IPSCs as DNQX blocked both EPSCs and IPSCs, we also observed monosynaptic IPSCs. Both EPSC amplitude (P=0.02) and IPSC amplitude (P=0.02) were significantly higher in female mice [14 neurons (7 female mice) and 16 neurons (6 male mice)]. The E/l ratio did not show a significant sex difference (P=0.14). The left panel shows an example of monosynaptic EPSC and IPSC in an SST neuron. 4-AP rescues TTX-induced suppression of EPSC and IPSC, the monosynaptic nature of which is revealed in the presence of AMPAand NMDAreceptor blockers (DNQX+AP5). The addition of bicuculline confirmed the GABAergic nature of IPSCs. Scale 50pA/5ms. C) EPSC amplitude, IPSC amplitude, and E/l ratio in PL-mPFC (pyramidal) neurons in response to light activation of PL-mPFC (VGLUT3) neurons. We observed monosynaptic EPSCs in pyramidal neurons. Both monosynaptic and disynaptic IPSCs were present in pyramidal neurons. Neither EPSC amplitude (P=0.28) nor IPSC amplitude (P=0.36) showed a significant sex difference [12 neurons (5 female mice) and 13 neurons (5 male mice)]. However, the E/l ratio showed a significant sex difference (P=0.04). The left panel shows an example of monosynaptic EPSC and IPSC in a pyramidal neuron. 4-AP rescues TTX-induced suppression of EPSC and IPSC, the monosynaptic nature of which is revealed in the presence of AMPA and NMDA receptor blockers (DNQX+AP5). The addition of bicuculline confirmed the GABAergic nature of IPSCs. Scale 50pA/5ms. Blue line indicateslight application.

## Discussion

Our studies show MRN (VGLUT3) neuron-mediated monosynaptic activation of mPFC (VGLUT3) neurons. The MRN (VGLUT3) neuron-mediated polysynaptic responses in mPFC VGLUT3 and SST neurons but not in PV or pyramidal neurons suggest the possibility that MRN (VGLUT3) neurons preferentially innervate mPFC (VGLUT3) neurons that form synapses onto local SST neurons and VGLUT3 neurons. Consistent with MRN (VGLUT3) neuron-mediated excitatory responses in these mPFC GABAergic neurons, activation of MRN (VGLUT3) terminals produced a long-lasting enhancement of GABAergic transmission and a long-lasting depression of excitatory synaptic transmission in PL-mPFC pyramidal neurons from female but not male mice. In agreement with MRN (VGLUT3) neuron-mediated plasticity in the PL-mPFC, we observed a suppression of fear memory in female but not male mice following the chemogenetic activation of MRN (VGLUT3) neurons. These results are significant given the role of the PL-mPFC in fear memory expression (*1, 29, 30*).

MRN neurons are functionally heterogeneous. In addition to GABAergic neurons, the MRN contains neurons expressing VGLUT2, VGLUT3, and 5-HT (*14–17*). Most MRN neurons projecting to the mPFC are VGLUT3-expressing neurons (*10*). An earlier study showed that raphe serotonergic neurons activate mPFC GABAergic neurons by releasing glutamate (*19*). Also, the VGLUT3-expressing raphe axons co-localize with cortical VGLUT3 neurons (*23*). Our experiments demonstrate that MRN (VGLUT3) neurons activate VGLUT3 and SST neurons in the mPFC. This neuromodulation is particularly significant as the dual release of glutamate and GABA from mPFC (VGLUT3) neurons, and SST neuron-mediated GABA release affect pyramidal neuron activity (*25, 31, 32*). Consistently, a 10Hz light activation of MRN (VGLUT3) terminals produces; 1) hyperpolarization, 2) a long-lasting enhancement of GABAergic transmission, and 3) a long-lasting suppression of glutamatergic transmission in mPFC pyramidal neurons. Therefore, MRN (VGLUT3) neuron-mediated activation of mPFC VGLUT3 and SST neurons and the resulting GABAergic inhibition of mPFC pyramidal neuron activity might play a role in the suppression of fear memory in female mice.

The female-specific mPFC plasticity and fear memory suppression might be due to a combined effect of multiple synapses along the MRN (VGLUT3)-mPFC pathway. The basal glutamatergic but not GABAergic transmission in MRN (VGLUT3) neurons is diminished in female mice compared to male mice. Furthermore, fear conditioning involves an enhanced glutamatergic transmission in the MRN (VGLUT3) neurons of female mice but not male mice. A diminished basal glutamatergic transmission in MRN (VGLUT3) neurons and its potentiation in response to fear conditioning might be a critical mechanism for MRN (VGLUT3)-mediated suppression of fear memory in female mice. This experience-dependent glutamatergic plasticity might make MRN (VGLUT3) neurons more amenable to chemogenetic activation, leading to an attenuation of PL-mPFC function and hence the suppression of fear memory in female mice. Also, the higher basal glutamatergic transmission in MRN (VGLUT3) neurons in male mice might result in an increased bottom-up modulation of the mPFC, and this effect might occlude the impact of chemogenetic activation. The diminished glutamatergic drive onto MRN (VGLUT3) neurons in female mice compared to male mice could contribute to the sex difference in behaviors relevant to anxiety- or psychological trauma-related disorders (*33–35*).

In addition to the sex difference in glutamatergic transmission and plasticity in MRN (VGLUT3) neurons, the PL-mPFC (VGLUT3)-SST neuron transmission is significantly higher in females compared to males. Also, the E/I ratio of the PL-mPFC (VGLUT3) neuron-pyramidal neuron transmission in female mice is lower than that of male mice. Given that MRN (VGLUT3) neurons primarily target VGLUT3 neurons in the mPFC, sex difference in PL-mPFC (VGLUT3) neuron-mediated transmission in local SST and pyramidal neurons might be an additional mechanism that augments the effect of MRN (VGLUT3) neuron-mediated inhibition of PL-mPFC pyramidal neurons in female mice. Thus, the sex difference in both the synaptic transmission in MRN (VGLUT3) neurons and the synaptic output of PL-mPFC (VGLUT3) neurons might be responsible for female-specific synaptic plasticity in the PL-mPFC and fear memory suppression following the activation of MRN (VGLUT3) neurons. Chromosomal and hormonal effects might play a role in establishing these sex-specific circuits (*36*).

Although not addressed in the current study, the release of 5-HT from MRN (VGLUT3) neurons could contribute to the observed synaptic and behavioral effects (*10, 19, 37*). This potential role of 5-HT is particularly important as VGLUT3/CCK GABAergic neurons express ionotropic 5-HT3a receptors (*38–39*). Furthermore, other 5-HT3a receptor-expressing GABAergic neurons including vasoactive intestinal polypeptide neurons might receive synaptic inputs from MRN (VGLUT3) neurons (*38*). Therefore, future studies will be necessary to determine the role of 5-HT release from MRN (VGLU3) neurons in both mPFC function and the regulation of fear behavior.

In conclusion, our study demonstrates that MRN (VGLUT3) neurons modulate the mPFC through glutamatergic transmission in VGLUT3 and SST GABAergic neurons. Furthermore, this study describes sex-specific synaptic transmission and plasticity in the MRN (VGLUT3)-mPFC neuromodulatory pathway and their potential role in the suppression of fear memory in female mice. Given the increased prevalence of anxiety- and psychological trauma-related disorders in females compared to males, identifying synaptic and molecular targets in this neuromodulatory pathway that are responsive to pharmacological or behavioral interventions might provide critical insights into evidence-based approaches to managing psychiatric disorders.

## Materials and Methods

### Animals

Both female and male mice were used in all experiments. B6;129S-*Slc17a8^tm1.1(cre)Hze^*/J (Stock number: 028534, Jackson Laboratories) mice were used for studying the effect of optogenetic and chemogenetic activation experiments. We crossed B6;129S-*Slc17a8^tm1.1(cre)Hze^*/J mice with GIN mice (FVB-Tg(GadGFP)45704Swn/J, Stock number: 003718, Jackson Laboratories) and G42 mice (CB6-Tg(Gad1-EGFP)G42Zjh/J, Stock number: 007677, Jackson Laboratories) to generate mice that are positive for Cre in VGLUT3 neurons and EGFP in SST and PV neurons, respectively (*28–40*). VGLUT3-tdTomato mice were created by crossing B6;129S-*Slc17a8^tm1.1(cre)Hze^*/J with B6.Cg-*Gt(ROSA)26Sor^tm14(CAG-tdTomato)Hze^*/J (Stock number: 007914, Jackson Laboratories) (*41*). Mice were maintained on a 12-hour light-dark cycle at 23°C with access to food and water ad libitum. The Institutional Animal Care and Use Committee of the University of Toledo approved all the procedures.

### Stereotactic surgery

Viral injections were carried out on a stereotactic apparatus (Stoelting, IL, USA) under isoflurane anesthesia. 200 nl pAAV-EF1a-double floxed-hChR2(H134R)-mCherry-WPRE-HGHpA, pAAV-EF1a-double floxed-hChR2(H134R)-EYFP-WPRE-HGHpA, pAAV-hSyn-EGFP, pAAV-hSyn-DIO-hM3D(Gq)-mCherry or pAAV-hSyn-DIO-hM4D(Gi)-mCherry were injected at a rate of 100 nl/min using an injector (Nanoliter 2010, WPI, FL, USA) attached to a pulled glass injection micropipette (∼50 µm tip, Item no: 504949, WPI, FL, USA). Injection coordinates for MRN were (from bregma): AP: −3.6mm; ML: 1.2mm, DV: 4.6mm, 16° angle (ear bars at 18mm, anesthesia mask bar at 14mm). Injection coordinates for mPFC were (from bregma): AP: 2.1mm; ML: 0.8mm, DV: 2.6mm, 16° angle (ear bars at 18mm, anesthesia mask bar at 14mm). The accuracy of virus injections was confirmed using fluorescence microscopy to visualize mCherry/EGFP/EYFP-expressing neurons. Injected mice were used for electrophysiology and behavior experiments 3 weeks after the surgery.

### Electrophysiology

Following transcardial perfusion with ice-cold oxygenated artificial cerebrospinal fluid (ACSF) containing (in mM): NaCl (118), glucose (10), KCl (2.5), NaH_2_PO_4_ (1), CaCl_2_ (1) and MgSO_4_ (1.5) (325 mOsm, pH 7.4) under pentobarbital (120mg//kg) anesthesia, brains were isolated for preparing prefrontal cortical (300 µm) and MRN (200 µm) slices on a vibratome (Campden Instruments). Prefrontal cortical slices were incubated in ACSF at room temperature for a minimum of 1 hr, followed by the transfer of brain slices to the recording chamber superfused with the ACSF containing 2.5 mM CaCl_2_ at 32°C flowing at 2 ml/min rate. MRN slices were transferred to oxygenated ACSF at 35°C and allowed to recover for at least 1 hr, during which the ACSF cooled down to room temperature. Both the mPFC and MRN were located using a 4X objective. Recorded neurons were visualized using a 40X water immersion objective and video-enhanced differential interference contrast microscopy. EGFP- and tdTomato-expressing neurons were identified using fluorescence microscopy, while pyramidal neurons were identified by their morphology, location, and membrane characteristics. MRN (VLGUT3) neuron-mediated currents, the effect of 10Hz stimulation on EPSPs, and membrane properties in mPFC and MRN (VGLUT3) neurons were recorded using recording electrodes of 3–5 MΩ resistance filled with an internal solution containing (in mM): K-gluconate (130), KCl (10), MgCl_2_ (5), MgATP (5), NaGTP (0.2), HEPES (5), pH adjusted to 7.4 with KOH. EPSPs were evoked at 0.1 Hz in layer 5 pyramidal neurons by electrical stimulation (100-200 µA) of layer 2/3 using an extracellular stimulating electrode (FHC, ME, USA). Membrane properties in mPFC pyramidal neurons were recorded before and 25 min after the 10Hz light stimulation. The effect of 10Hz light stimulation on hyperpolarization and sIPSCs in pyramidal neurons was recorded using an internal solution containing (in mM): K-gluconate (130), KCl (5), MgCl_2_ (2), MgATP (5), NaGTP (0.2), HEPES (5), pH adjusted to 7.4 with KOH. sIPSCs were recorded in pyramidal neurons at 0 mV. These GABA_A_ receptor-mediated sIPSCs were confirmed using bicuculline (10 µM). mEPSCs and mIPSCs were recorded from MRN (VGLUT3) neurons at −50 mV and 0 mV, respectively, in the presence of tetrodotoxin (1 µM). AMPA receptor-mediated mEPSCs and GABA_A_ receptor-mediated mIPSCs were confirmed using DNQX (10 µM) and bicuculline (10 µM), respectively. An internal solution containing (in mM): Cesium methanesulfonate (130), NaCl (6), HEPES (5), MgATP (5), NaGTP (0.2), pH adjusted to 7.4 with CsOH was used for recording mEPSCs and mIPSCs in MRN (VGLUT3) neurons. mPFC (VGLUT3) neuron-mediated transmission in SST, PV, and pyramidal neurons were recorded at - 20mV using an internal solution containing (in mM): K-gluconate (130), KCl (2), MgCl_2_ (2), MgATP (5), NaGTP (0.2), HEPES (5), pH adjusted to 7.4 with KOH. Blue light (470 nm) was emitted from a Lumen 1600-LED (Prior) for optogenetic experiments. Electrophysiological recordings were rejected when series resistance or holding current changed by 10% or more.

### Behavior

A mouse was placed in the conditioning chamber (context A) within a soundproof box (Actimetrics, IL, USA) for fear conditioning. After a 2 min exploration, mice received 2 habituation tones (5 kHz, 30Db, 30 s duration) followed by 3 tones (30 s duration) that co-terminated with foot shocks (2 s duration, 0.5 mA, 30 s interval). Mice were returned to their home cages 30 s after the final tone-shock pairing. The control tone-alone (TA) group received tone presentations similar to the fear-conditioned (FC) group, except that they did not receive foot shocks. Fear-extinction (FE) training was carried out 24 hr after conditioning and consisted of 30 tone presentations (30 s duration) at 30 s intervals (context B). For all groups, fear memory was tested 48 hr after the initial fear conditioning/tone alone treatment by exposing the mice to 3 tones (30 s duration) at 30 s intervals (context B). Freezing was measured using FreezeFrame software (Actimetrics, IL, USA). For studying the effect of chemogenetic activation or suppression of MRN (VGLUT3) neurons on fear memory, fear-conditioned mice received either saline or CNO (1 mg/kg, intraperitoneal) 1 hr before fear extinction training. In control experiments to confirm the lack of a non-specific effect of CNO on fear memory and extinction, CNO or saline was administered to mice that received the pAAV-hSyn-EGFP injection into the MRN. For the experience-dependent plasticity experiments, brain slices were prepared within 4 hr after the fear memory test.

### Data analysis

We used Clampfit 10.7 (Molecular Devices, CA, USA) for analyzing synaptic responses and membrane properties. EPSP slope was measured at the initial 2 ms of EPSP response. Membrane properties were analyzed as described before, and frequency and accommodating properties of action potentials and shape, capacitance, and location of neurons were used in identifying pyramidal and SST neurons (*25, 42, 43*). Statistical analyses were performed using IBM SPSS statistics (version 27) or GraphPad Prism (version 9) software. A two-way ANOVA followed by the Bonferroni post hoc test or t-test was used for analyzing spontaneous currents, EPSPs, membrane properties, and behavioral data. Data from experience-dependent plasticity experiments were averaged per mouse before the statistical analysis.

### Funding

This work was supported by the National Institutes of Health (grant number HD076914 to I.N.).

## Acknowledgments

pAAV-EF1a-double floxed-hChR2(H134R)-mCherry-WPRE-HGHpA (20297-AAV9, Addgene) and pAAV-EF1a-double floxed-hChR2(H134R)-EYFP-WPRE-HGHpA (20298-AAV9, Addgene) were gifts from Dr. Karl Deisseroth. pAAV-hSyn-EGFP (50465-AAV2, Addgene), pAAV-hSyn-DIO-hM4D(Gi)-mCherry (44362-AAV2, Addgene) and pAAV-hSyn-DIO-hM3D(Gq)-mCherry (44361-AAV2, Addgene) were gifts from Dr. Bryan Roth. The authors thank Dr. Andrea Kalinoski (Integrated Core Facilities) for help with the experiments.

## Supplementary Figures and Legends

**Fig. S1.**
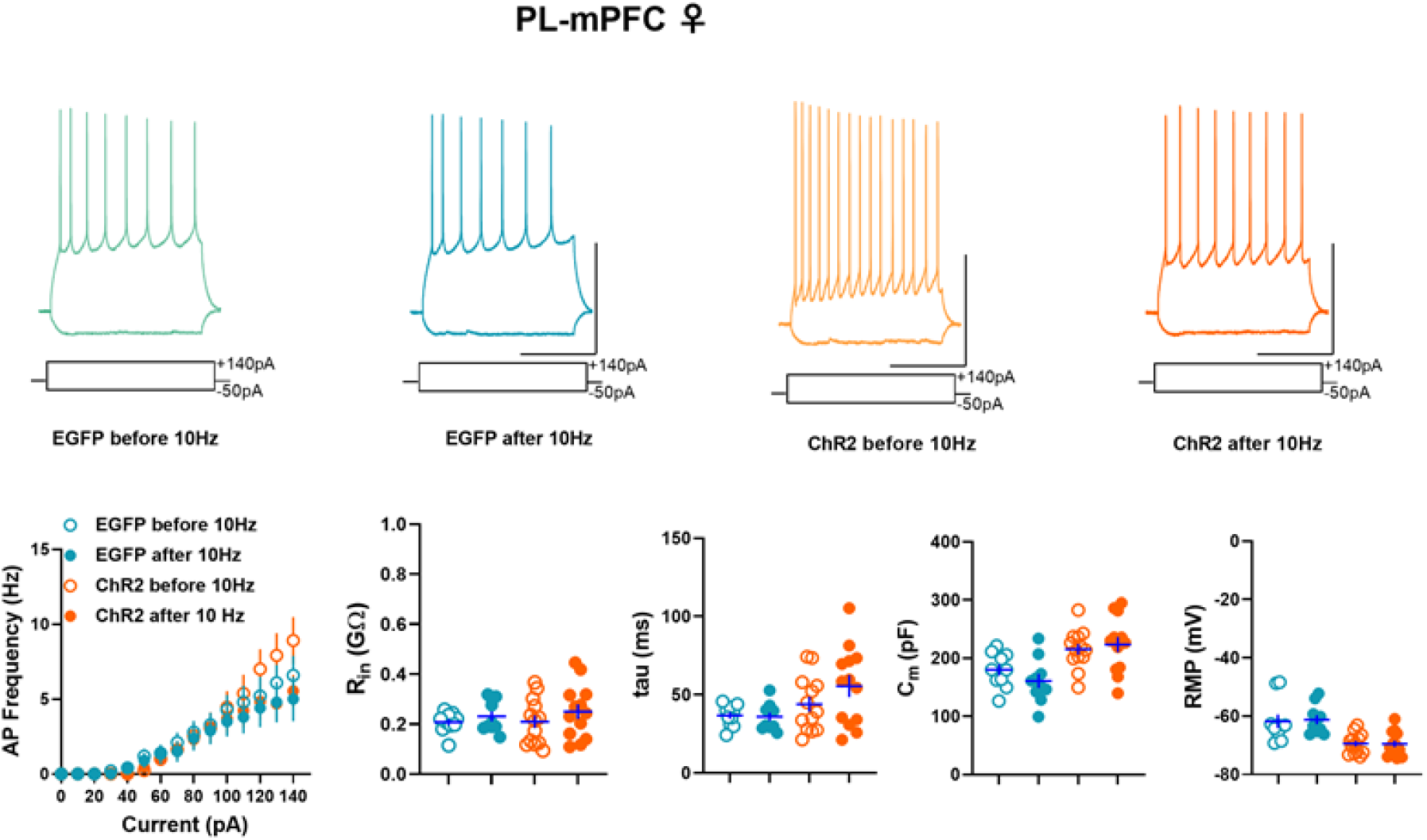
Effect of 10HZ light activation on active and passive membrane properties of PL-mPFC pyramidal neurons from female MRN (VGLUT3)-ChR2 (13 neurons/7 mice) and MRN-EGFP (10 neurons/5 mice) mice. Light activation did not affect the number of action potentials (F_3,42_=0 45, P=0.72), input resistance (Rin, F_3,42_=0.72, P=0.55), time constant (tau, F_3,42_=1.3, P=0.29), membrane capacitance (Cm, F_3,42_=2.58, P=0.07), or resting membrane potential (RMP, F_3,42_=2.07, P=0.12) in pyramidal neurons. The upper panel shows example traces before and after the 10Hz stimulation in PL-mPFC slices from MRN (VGLUT3)-ChR2 and MRN-EGFP mice. Scale: 50 mV/500 ms.

**Fig. S2.**
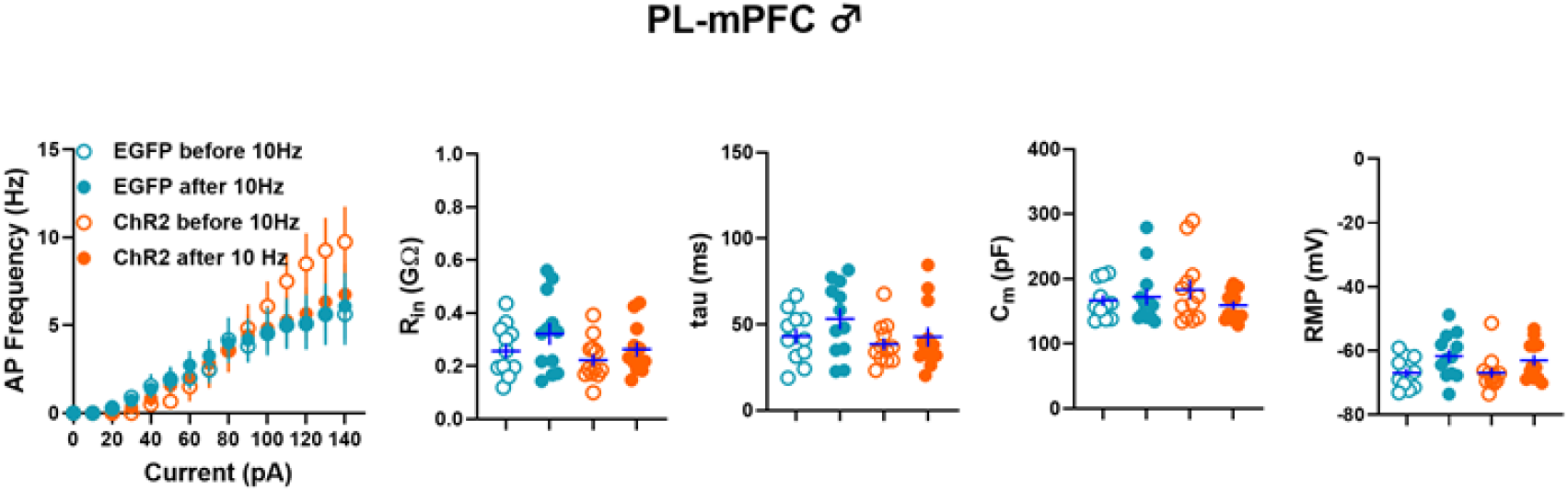
Effect of 10Hz light activation on active and passive membrane properties of PL-mPFC pyramidal neurons from male MRN (VGLUT3)-ChR2 (12 neurons/6 mice) and MRN-EGFP (12 neurons/6 mice) mice. Light activation did not affect the number of action potentials (F_3,44_=0.13, P=0.94), input resistance (Rin, F_3,44_=1.8, P=0.16), time constant (tau, F_3,44_=2.49, P=0.07), membrane capacitance (Cm, F_3,44_=0.82, P=0.48), an’d resting membrane potential (RMP, F_3,4_=2.4, P=0.08) in pyramidal neurons. •

**Fig. S3.**
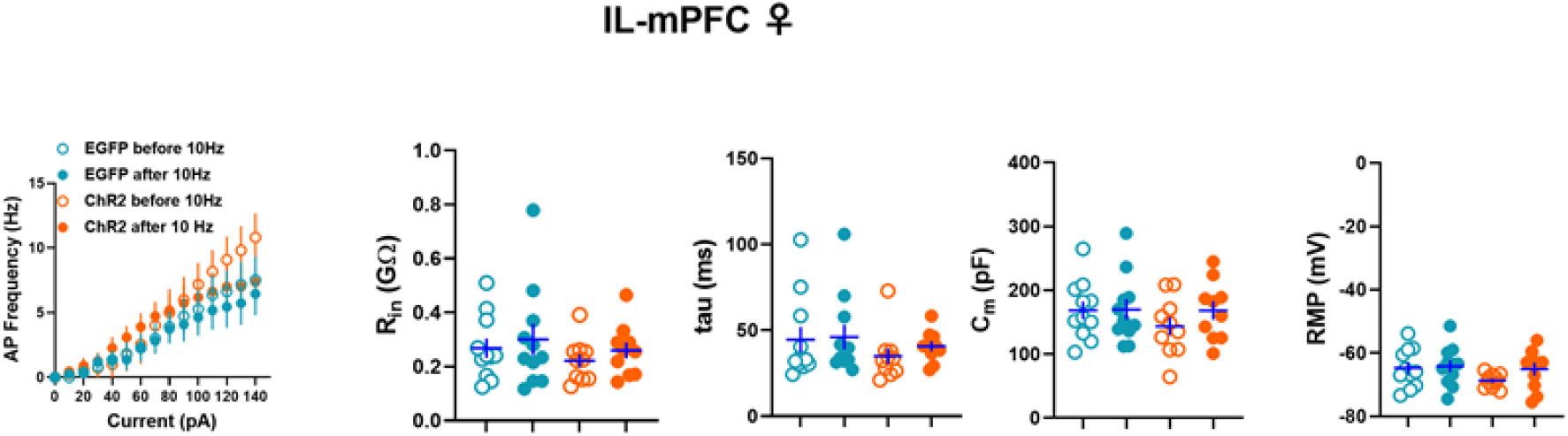
Effect of 10Hz light activation on active and passive membrane properties of IL-mPFC pyramidal neurons from female MRN (VGLUT3)-ChR2 (11 neurons/6 mice) and MRN-EGFP (11 neurons/5 mice) mice. Light activation did not affect the number of action potentials (F_3,40_=0.42, P=0.74), input resistance (Rin, F_3,40_=0.66, P=0.58), time constant (tau, F_3,40_=0.69, P=0.56), membra’ne capacitance (Cm, F_3,40_=0.71, P=0.55), or’resting membrane potential (RMP, F_3,40_=1.4, P=0.26) in pyramidal neurons.

**Fig. S4.**
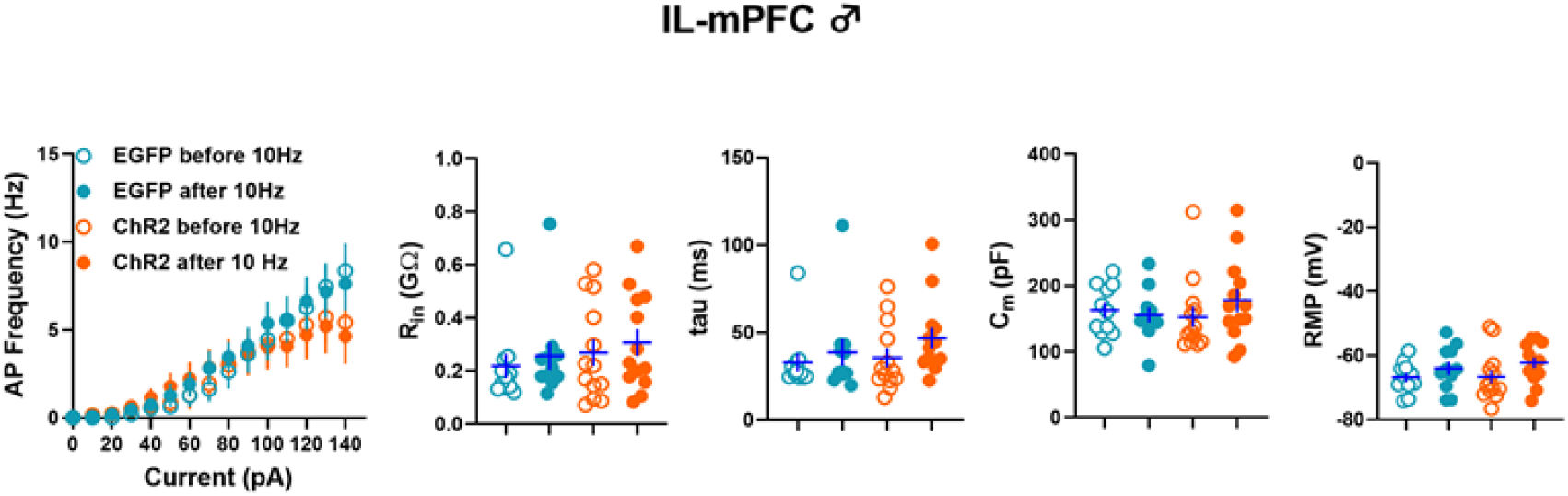
Effect of 10Hz light activation on active and passive membrane properties of IL-mPFC pyramidal neurons from male MRN (VGLUT3)-ChR2 (13 neurons/6 mice) and MRN-EGFP (11 neurons/5 mice) mice. Light activation did not affect the number of action potentials (F_3,44_=0.16, P=0.92), input resistance (Rin, (F_3,44_=0.52, P=0.67), time constant (tau, F_3,44_=1.01, P=0.4), membrane capacitance (Cm, F_3,44_=0.59, P=0.63), and resting membrane potential (RMP, F_3,44_;1.4, P=0.25) in pyramidal neurons. •

**Fig. S5.**
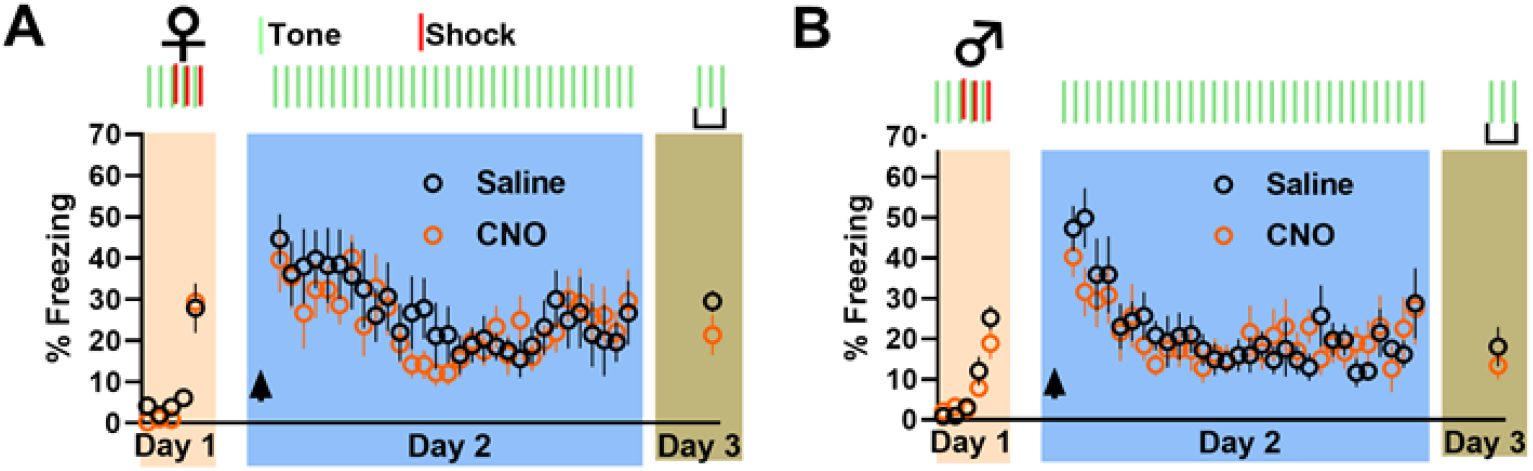
CNO (1 mg/kg)- and saline-treatment in female **(A)** [(CNO,n=10, saline, n=10), (F_1,18_=0.17, P=0.68)] and male **(B)** [(CNO,n=9, saline, n=9), (F_1,16_=0.2, P=0.8)] C57BL6/J mice that received a pAAV-hSyn-EGFP injection into the MRN did not show a significant difference in fear memory or extinction.

**Fig. S6.**
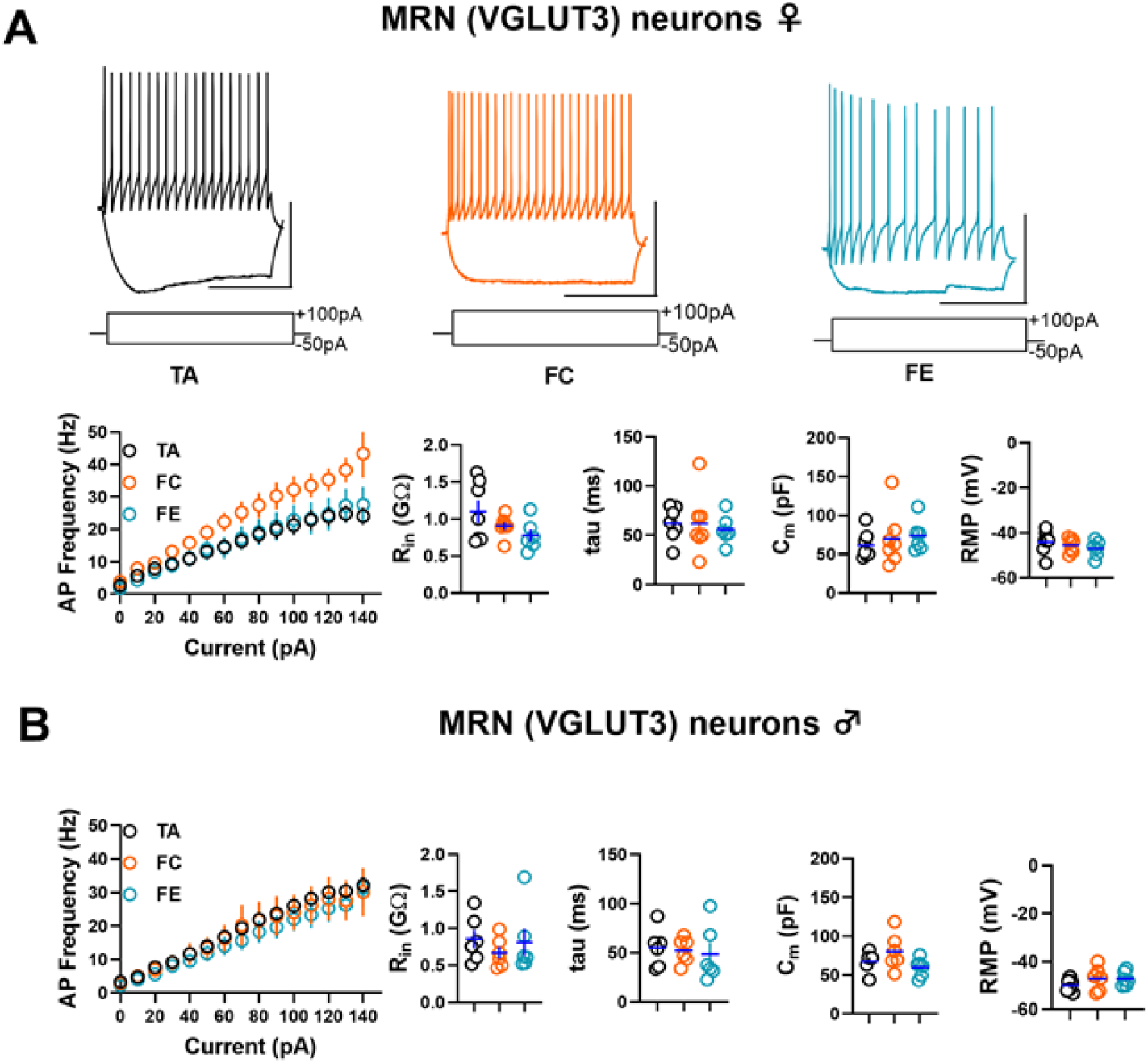
Effect of fear conditioning and extinction on the excitability of MRN (VGLUT3) neurons. A) Passive and active membrane properties in tdTomato-expressing MRN (VGLUT3) neurons from female tone-alone (TA, 20 neurons/7 mice), fear-condition (FC, 16 neurons/7 mice), and fear-extinction (FE, 15 neurons/6 mice) groups. We did not observe a significant change in the number of action potentials (F_2,17_=2.7, P=0.09), input resistance (Rin, F_2,17_=2.23, P=0.14), time constant (tau, F_2,17_=0.15, P=0.85), membrane capacitance (Cm, F_2,17_=0.38, P=0.69), and resting membrane’potential (RMP, F_2,17_=0.8, P=0.46) among TA, FC, and FE groups. The upper panel shows example traces. Scale: 50 mV/500 ms. B) We did not observe a significant change in the number of action potentials (F_2,15_=0.2, P=0.82), input resistance (Rin, F_2,15_=0.49, P=0.62), time constant (tau, F_2,15_=0.14, P =0.87), membrane capacitance (Cm, F_2,15_ =2.3, P=0.13), and resting membrane’ potential (RMP, F_2,15_=1.2, P=0.33) among male TA (17 neurons/6 mice), FC (16 neurons/6 mice) and FE (15 ne’urons/6 mice) groups. A comparison of membrane properties in control TA groups did not show a difference in the number of action potentials (F_1,11_=1.1, P=0.3), input resistance (P=0.25), time constant (P=0.49) or capacitance (P=0.52) b etween female and male groups. However, the RMP comparison showed a significant depolarization in the female TA group compared to the male TA group (P=0.02).

**Fig. S7.**
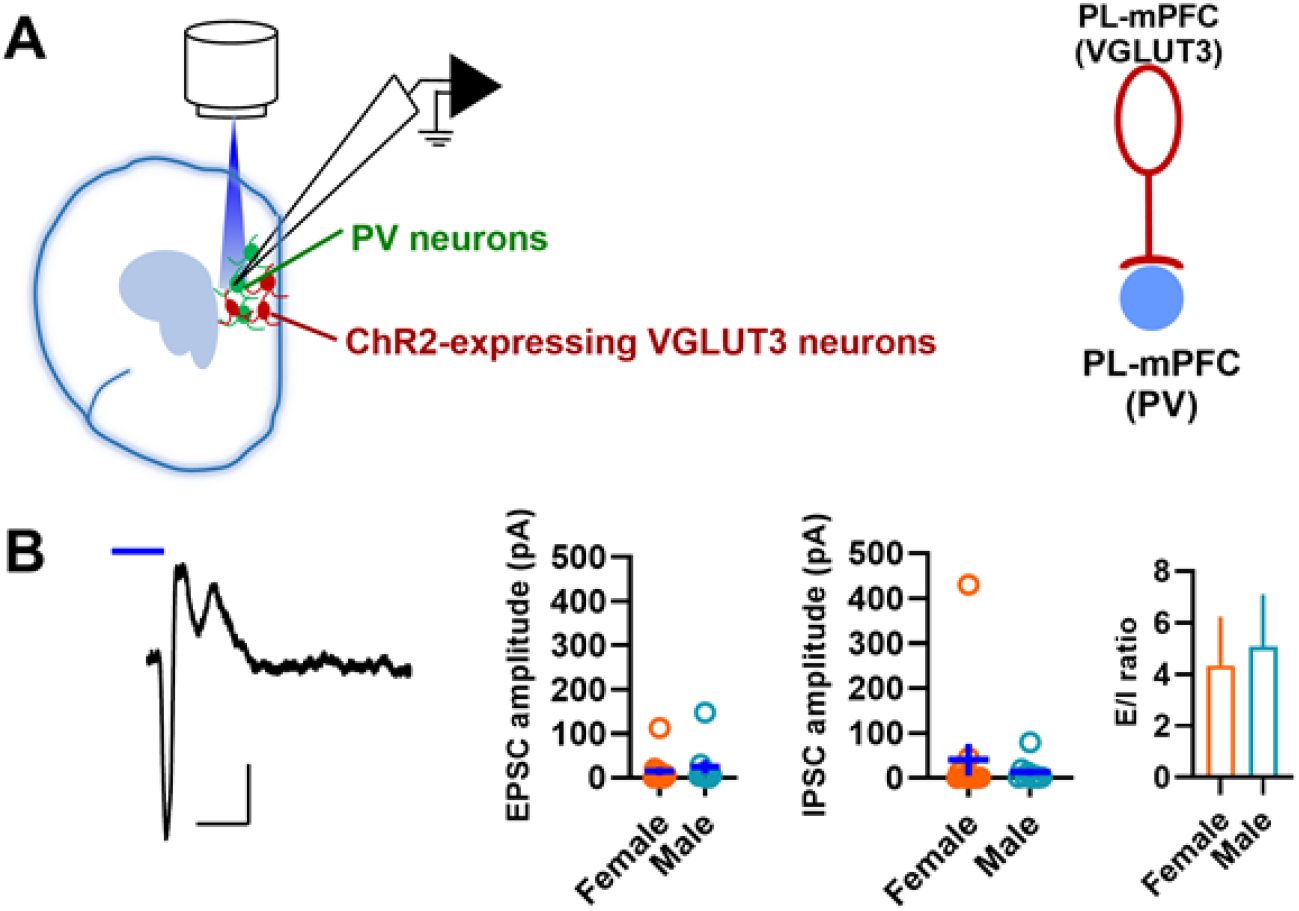
PL-mPFC (VGLUT3) neuron-mediated glutamatergic and GABAergic currents in local PV neurons. A) schematic of light activation of PL-mPFC (VGLUT3) neurons and whole-cell recording in local PV neurons. B) EPSC amplitude, IPSC amplitude, and E/1 ratio in PL-mPFC (PV) neurons in response to light activation of PL-mPFC (VGLUT3) neurons. We did not observe sex difference in EPSC amplitude (P=0.57), IPSC amplitude (P=0.52), or E/1 ratio (P=0.79) [12 neurons (3 female mice) and 9 neurons (3 male mice)]. The left panel shows an example of glutamatergic and GABAergic currents in PV neurons in response to light activation of mPFC (VGLUT3) neurons. Scale 50pN5ms. Blue line indicates light application.

## References

1. K. A. Corcoran, G. J. Quirk, Activity in prelimbic cortex is necessary for the expression of learned, but not innate, fears. J Neurosci 27, 840–844 (2007).

2. M. R. Milad, G. J. Quirk, Neurons in medial prefrontal cortex signal memory for fear extinction. Nature 420, 70–74 (2002).

3. B. M. Graham, M. R. Milad, The study of fear extinction: implications for anxiety disorders. Am J Psychiatry 168, 1255–1265 (2011).

4. R. M. Guthrie, R. A. Bryant, Extinction learning before trauma and subsequent posttraumatic stress. Psychosom Med 68, 307–311 (2006).

5. M. Wicking et al., Deficient fear extinction memory in posttraumatic stress disorder. Neurobiol Learn Mem 136, 116–126 (2016).

6. D. G. Balazsfi et al., Median raphe region stimulation alone generates remote, but not recent fear memory traces. PLoS One 12, e0181264 (2017).

7. V. Avanzi, V. M. Castilho, T. G. de Andrade, M. L. Brandao, Regulation of contextual conditioning by the median raphe nucleus. Brain Res 790, 178–184 (1998).

8. T. G. Andrade, H. Zangrossi, Jr., F. G. Graeff, The median raphe nucleus in anxiety revisited. J Psychopharmacol 27, 1107–1115 (2013).

9. R. C. Almada, K. G. Borelli, L. Albrechet-Souza, M. L. Brandão, Serotonergic mechanisms of the median raphe nucleus-dorsal hippocampus in conditioned fear: Output circuit involves the prefrontal cortex and amygdala. Behavioural brain research 203, 279–287 (2009).

10. A. Szonyi et al., The ascending median raphe projections are mainly glutamatergic in the mouse forebrain. Brain Struct Funct 221, 735–751 (2016).

11. E. G. Meloni, C. L. Reedy, B. M. Cohen, W. A. Carlezon, Jr., Activation of raphe efferents to the medial prefrontal cortex by corticotropin-releasing factor: correlation with anxiety-like behavior. Biological psychiatry 63, 832–839 (2008).

12. R. P. Vertes, W. J. Fortin, A. M. Crane, Projections of the median raphe nucleus in the rat. The Journal of comparative neurology 407, 555–582 (1999).

13. A. Teissier et al., Activity of Raphé Serotonergic Neurons Controls Emotional Behaviors. Cell Rep 13, 1965–1976 (2015).

14. K. E. Sos et al., Cellular architecture and transmitter phenotypes of neurons of the mouse median raphe region. Brain Struct Funct 222, 287–299 (2017).

15. A. Szőnyi et al., Median raphe controls acquisition of negative experience in the mouse. Science 366, eaay8746 (2019).

16. B. W. Okaty, K. G. Commons, S. M. Dymecki, Embracing diversity in the 5-HT neuronal system. Nat Rev Neurosci 20, 397–424 (2019).

17. C. Gras et al., A third vesicular glutamate transporter expressed by cholinergic and serotoninergic neurons. The Journal of neuroscience : the official journal of the Society for Neuroscience 22, 5442–5451 (2002).

18. M. Hajós, S. E. Gartside, V. Varga, T. Sharp, In vivo inhibition of neuronal activity in the rat ventromedial prefrontal cortex by midbrain-raphe nuclei: role of 5-HT1A receptors. Neuropharmacology 45, 72–81 (2003).

19. Q. Sun et al., A whole-brain map of long-range inputs to GABAergic interneurons in the mouse medial prefrontal cortex. Nat Neurosci 22, 1357–1370 (2019).

20. J. Somogyi et al., GABAergic basket cells expressing cholecystokinin contain vesicular glutamate transporter type 3 (VGLUT3) in their synaptic terminals in hippocampus and isocortex of the rat. Eur J Neurosci 19, 552–569 (2004).

21. C. Fasano et al., Regulation of the Hippocampal Network by VGLUT3-Positive CCK-GABAergic Basket Cells. Front Cell Neurosci 11, 140 (2017).

22. K. A. Pelkey et al., Paradoxical network excitation by glutamate release from VGluT3(+) GABAergic interneurons. Elife 9, (2020).

23. R. A. Senft, M. E. Freret, N. Sturrock, S. M. Dymecki, Neurochemically and Hodologically Distinct Ascending VGLUT3 versus Serotonin Subsystems Comprise the r2-Pet1 Median Raphe. J Neurosci 41, 2581–2600 (2021).

24. L. Petreanu, T. Mao, S. M. Sternson, K. Svoboda, The subcellular organization of neocortical excitatory connections. Nature 457, 1142–1145 (2009).

25. P. Koppensteiner et al., Diminished Fear Extinction in Adolescents Is Associated With an Altered Somatostatin Interneuron-Mediated Inhibition in the Infralimbic Cortex. Biol Psychiatry 86, 682–692 (2019).

26. A. A. Oliva, Jr., M. Jiang, T. Lam, K. L. Smith, J. W. Swann, Novel hippocampal interneuronal subtypes identified using transgenic mice that express green fluorescent protein in GABAergic interneurons. The Journal of neuroscience : the official journal of the Society for Neuroscience 20, 3354–3368 (2000).

27. T. L. Daigle et al., A Suite of Transgenic Driver and Reporter Mouse Lines with Enhanced Brain-Cell-Type Targeting and Functionality. Cell 174, 465–480.e422 (2018).

28. B. Chattopadhyaya et al., Experience and activity-dependent maturation of perisomatic GABAergic innervation in primary visual cortex during a postnatal critical period. J Neurosci 24, 9598–9611 (2004).

29. I. Vidal-Gonzalez, B. Vidal-Gonzalez, S. L. Rauch, G. J. Quirk, Microstimulation reveals opposing influences of prelimbic and infralimbic cortex on the expression of conditioned fear. Learn Mem 13, 728–733 (2006).

30. T. F. Giustino, S. Maren, The Role of the Medial Prefrontal Cortex in the Conditioning and Extinction of Fear. Front Behav Neurosci 9, 298 (2015).

31. A. Adler, R. Zhao, M. E. Shin, R. Yasuda, W. B. Gan, Somatostatin-Expressing Interneurons Enable and Maintain Learning-Dependent Sequential Activation of Pyramidal Neurons. Neuron 102, 202–216.e207 (2019).

32. J. Urban-Ciecko, E. E. Fanselow, A. L. Barth, Neocortical somatostatin neurons reversibly silence excitatory transmission via GABAb receptors. Curr Biol 25, 722–731 (2015).

33. G. E. Fenton, D. M. Halliday, R. Mason, T. W. Bredy, C. W. Stevenson, Sex differences in learned fear expression and extinction involve altered gamma oscillations in medial prefrontal cortex. Neurobiol Learn Mem 135, 66–72 (2016).

34. S. Börchers, J. P. Krieger, M. Asker, I. Maric, K. P. Skibicka, Commonly-used rodent tests of anxiety-like behavior lack predictive validity for human sex differences. Psychoneuroendocrinology 141, 105733 (2022).

35. J. J. Botterill et al., Bidirectional Regulation of Cognitive and Anxiety-like Behaviors by Dentate Gyrus Mossy Cells in Male and Female Mice. J Neurosci 41, 2475–2495 (2021).

36. E. Lentini, M. Kasahara, S. Arver, I. Savic, Sex differences in the human brain and the impact of sex chromosomes and sex hormones. Cereb Cortex 23, 2322–2336 (2013).

37. P. Celada, M. V. Puig, F. Artigas, Serotonin modulation of cortical neurons and networks. Front Integr Neurosci 7, 25 (2013).

38. Z. J. Huang, A. Paul, The diversity of GABAergic neurons and neural communication elements. Nat Rev Neurosci 20, 563–572 (2019).

39. M. Morales, F. E. Bloom, The 5-HT3 receptor is present in different subpopulations of GABAergic neurons in the rat telencephalon. J Neurosci 17, 3157–3167 (1997).

40. A. A. Oliva, Jr., M. Jiang, T. Lam, K. L. Smith, J. W. Swann, Novel hippocampal interneuronal subtypes identified using transgenic mice that express green fluorescent protein in GABAergic interneurons. J Neurosci 20, 3354–3368 (2000).

41. L. Madisen et al., A robust and high-throughput Cre reporting and characterization system for the whole mouse brain. Nat Neurosci 13, 133–140 (2010).

42. S. A. Collins, Ninan, I., Development-dependent plasticity in vasoactive intestinal polypeptide neurons in the infralimbic cortex. Cerebral Cortex Communications 2, tgab007 (2021).

43. P. Koppensteiner, C. Galvin, I. Ninan, Lack of experience-dependent intrinsic plasticity in the adolescent infralimbic medial prefrontal cortex. Synapse 73, e22090 (2019).

